# Novel RNA viruses reveal a complex mycovirome in the smut fungus *Thecaphora thlaspeos*

**DOI:** 10.64898/2026.03.24.713915

**Authors:** Andres G. Jacquat, Nicolas Bejerman, Humberto Debat

## Abstract

Mycoviruses are widespread in fungi, yet their diversity and host associations remain poorly explored in many lineages, including smut fungi. Here, we report the discovery of the first eight novel RNA viruses infecting the Brassicaceae-associated smut fungus *Thecaphora thlaspeos*. Using transcriptomic datasets derived from fungal mycelium and host plant controls, we identified dsRNA viral genomes supported by consistent read abundance, high genome coverage, and codon usage patterns closely matching those of the fungal host. All genomes are monosegmented and bicistronic, encoding capsid protein and polymerase genes in compact arrangements. Genome annotation and phylogenetic analyses based on the predicted RNA polymerase classified these viruses within the genera *Totivirus* and *Eimeriavirus*. Comparative analyses across fungal strains revealed intraspecific variation in virome composition, suggesting that host genetic background may influence viral community structure and that multiple dsRNA infection is common. Together, these findings expand the known diversity of mycoviruses in Ustilaginomycotina and identify *T. thlaspeos* as a host of a complex RNA virome. This work establishes a foundation for future studies on virus prevalence, transmission, and potential impacts on fungal biology and plant pathogenesis.

## Introduction

Mycoviruses, or fungal viruses, are ubiquitous infectious agents that inhabit fungi across major taxonomic groups, including plant pathogens, endophytes, and saprotrophs. They are currently classified into 47 viral families recognized by the International Committee on Taxonomy of Viruses (ICTV, https://ictv.global/vmr). Most mycoviruses possess double-stranded RNA (dsRNA) or positive-sense single-stranded RNA (+ssRNA) genomes and lack an extracellular transmission phase, instead relying on intracellular transfer via hyphal anastomosis and vertical transmission through spores (**Ghabrial et al., 2015; Hough et al., 2023**). While many infections are persistent or cryptic, accumulating evidence demonstrates their capacity to modulate fungal phenotypes, affecting vegetative growth, sporulation, and secondary metabolism (**Ghabrial et al., 2015**). Notably, virus-mediated hypovirulence in phytopathogenic systems has positioned mycoviruses as promising candidates for biological control strategies (**García-Pedrajas et al., 2019**).

The advent of high-throughput sequencing (HTS) has significantly expanded our understanding of fungal viromes, revealing complex assemblages of RNA viruses and frequently uncovering novel taxa (**Sutela et al., 2020; Jo et al., 2022; Nibert et al., 2019; Debat et al., 2025a**). Mining fungal transcriptomes for viral signatures has demonstrated that multi-virus coinfection is common and that virome composition can vary substantially among strains of the same fungal species. Large-scale transcriptome mining studies have identified complex mycoviromes comprising both dominant and low-abundance viruses across diverse fungal taxa (**Jo et al., 2022; Liu et al., 2025**). For example, analyses of multiple strains of *Corynespora cassiicola*, showed that each strain harbored distinct combinations of viruses, with individual isolates carrying between one and nine viral contigs belonging to different viral families. This highlights extensive strain-level variability and frequent multi-virus coinfections within a single fungal species (**Liu et al., 2025**). In the rice false smut pathogen *Ustilaginoidea virens*, dsRNA screening of 198 isolates revealed that approximately 95% harbored mycoviruses, predominantly members of the families *Totiviridae* and *Partitiviridae*, with mixed infections being the norm rather than the exception (**Jiang et al., 2015**). Such high prevalence and diversity of mycoviral infections within smut-related fungi raise the question of whether analogous viral communities exist in other, less-studied members of the *Ustilaginomycotina*.

Species of the genus *Thecaphora* include important plant pathogens of dicotyledonous hosts, such as the peanut smut fungus *Thecaphora frezii*. Although maize smut fungus *Ustilago maydis* have served as classical models for studying plant–fungus interactions, comparatively little is known about the biology of *Thecaphora* species despite their agronomic relevance (**Arias et al., 2021**). *Thecaphora thlaspeos* is a biotrophic basidiomycete and the only known smut fungus adapted to Brassicaceae hosts (**Frantzeskakis et al., 2017**), which are considered a critical model for both fundamental and applied plant pathology due to their susceptibility to yield and quality reductions (**Kronstad, 1996; Frantzeskakis et al., 2017; Plücker et al., 2021**). *T. thlaspeos* forms systemic infections in both roots and aerial tissues of species such as *Arabis hirsuta* and *Arabis alpina*, and it can also colonize the model plant *Arabidopsis thaliana* (**Frantzeskakis et al., 2017**). Unlike the well-characterized grass smut *U. maydis*, *T. thlaspeos* exhibits distinctive biological features, including plant signal-dependent teliospore germination, filamentous haploid growth, and the ability to overwinter with perennial hosts (**Courville et al., 2019**). The genome and transcriptome of *T. thlaspeos* have been characterized, revealing a typical smut genome architecture with both conserved and novel effector proteins (**Courville et al., 2019**). Our analyses of currently available genomic and transcriptomic datasets for *T. frezii* (**Arias et al., 2025**) have not revealed detectable viral sequence signatures. This absence prompted us to explore the virome of *T. thlaspeos*, whose publicly available RNA-seq datasets provide an opportunity for systematic transcriptome mining, and the potential to establish *T. thlaspeos* as a model system for investigating virus-host interactions within *Thecaphora* smut fungi.

In this work, we describe the discovery and characterization of eight novel RNA viruses in *T. thlaspeos*, identified through the reanalysis of RNA-seq data originally generated by **Courville *et al*. (2019).** The virome includes members of the genus *Eimeriavirus* (family *Pseudototiviridae*) and the genus *Totivirus* (family *Orthototiviridae*), the latter being one of the best-characterized mycoviral lineages (**Wickner et al., 2013**). We provide evidence for the fungal specificity of all eight viruses, document strain-level differences in virome composition between two *T. thlaspeos* mating types, and discuss these findings in the context of current knowledge of mycoviral ecology and smut fungus biology.

## Materials and methods

### Data source and virus discovery

RNA-seq datasets used in this study were retrieved from the Sequence Read Archive (SRA) and correspond to the genomic and transcriptomic work on *T. thlaspeos* by **Courville et al. (2019)**. Thirteen RNA libraries linked to the BioProject PRJEB24478 were analyzed: five derived from the smut fungus *T. thlaspeos* and eight from the host plant *Arabis hirsuta*. Four libraries were biological replicates of *T. thlaspeos* strain LF2 (mating type a2b2; samples THTG_LF21–THTG_LF24; runs ERR2305020–ERR2305023) and were sequenced in paired-end mode with an average read length of ∼270 bp. One additional library corresponded to *T. thlaspeos* strain LF1 (mating type a1b1; sample THTG_LF11; run ERR2305652) and was sequenced in single-end mode with 151 bp reads. The remaining eight libraries originated from *A. hirsuta* tissues and served as plant controls (samples ARHIR1–ARHIR4 and THTG_AH1–THTG_AH4; runs ERR2305034–ERR2305041); these paired-end libraries (average read length ∼270 bp) enabled discrimination between fungal-associated viral sequences and potential plant-associated viruses. Library sizes ranged from 12.9 to 42.2 million reads.

Direct quantitative comparison between strains LF2 and LF1 is constrained by several factors, including the absence of biological replicates for LF1, the lack of three viruses in that strain, and differences in sequencing strategies (paired-end versus single-end reads with different read lengths). Consequently, most comparative and quantitative analyses of viral relative abundance and genetic divergence were conducted using the LF2 transcriptomic dataset.

Virus discovery was implemented as previously described (**Debat et al., 2024; Debat et al., 2025b; Bejerman et al., 2025**). Briefly, raw nucleotide sequence reads from 13 RNA-seq libraries (five fungal mycelium and eight *Arabis hirsuta* plant control libraries) were downloaded from the NCBI Sequence Read Archive (SRA) under BioProject PRJEB24321. Datasets were pre-processed by trimming and filtering with the Trimmomatic v0.40 tool (http://www.usadellab.org/cms/?page=trimmomatic; acceded November 20, 2025) using standard parameters, except that the minimum average quality threshold was raised from 20 to 30 (initial ILLUMINACLIP step, sliding window trimming, average quality required = 30). The filtered reads were assembled *de novo* using rnaSPAdes with standard parameters on the Galaxy server (https://usegalaxy.org/; acceded November 20, 2025). The resulting transcripts were subjected to bulk local BLASTX searches (E-value < 1 × 10⁻⁵) against RefSeq protein sequences available at NCBI (https://www.ncbi.nlm.nih.gov/protein/?term=txid10239[Organism:exp]). Viral sequence hits were examined in detail. Tentative virus-like contigs were curated through iterative mapping of filtered reads from each SRA library; in each round, a subset of reads corresponding to the contig of interest was extracted and used to extend the sequence, which was then employed as the query for the next iteration. This process was repeated until no further extension was possible. The final extended and polished transcripts were reassembled using the alignment tool of Geneious v8.1.9 (Biomatters Ltd.) with high sensitivity parameters.

### Functional, structural and quantitative genomic analysis

Reads from each library were mapped to the reconstructed virus reference genomes, and raw read counts were recorded per virus per sample using Bowtie2 (**Langmead and Salzberg, 2012**) algorithm with default parameters. Two complementary abundance metrics were computed for each virus in each library. The reads per kilobase per million mapped reads (RPKM) was calculated as: RPKM = (read count × 10⁹) / (total library reads × genome size in nt). This metric normalizes for both sequencing depth and genome length, thereby enabling cross-sample and cross-virus comparisons. Mean genome coverage was calculated as: coverage = (read count × average read length) / genome size in nt, providing an intuitive measure of how many times the virus genome is covered on average by sequencing reads. For strain LF2, mean and standard deviation were computed across the four biological replicates. The coefficient of variation (CV = SD/mean × 100) was used to assess replicate consistency.

DNA sequencing libraries from the same bioproject (PRJEB24478) were analyzed to confirm the RNA nature of the discovered viral sequences. Eight paired-end DNA libraries generated for *T. thlaspeos* genome assembly (ERR2305024-ERR2305031) were mapped against all eight viral genome assemblies using Bowtie2 with default parameters. Raw DNA sequencing reads were obtained from the European Nucleotide Archive and processed without quality trimming to maintain maximum sensitivity for potential viral sequence detection.

Pairwise sequence identities were computed by direct positional comparison of MAFFT v7.511 (**Katoh et al., 2002**) aligned sequences. For each pair, the number of matching positions across the shorter of the two sequences was divided by the length of the longer sequence and expressed as a percentage. The standard genetic code was used for translation via Biopython’s 1.86 Bio.Seq.translate. Relative Synonymous Codon Usage (RSCU) was calculated following the standard formula: RSCU_ij = (X_ij × n_i) / Σ X_ij, where X_ij is the observed count of codon j for amino acid i, and n_i is the number of synonymous codons encoding amino acid i. Codon counts were tallied over the coding genome sequence read in frame (triplets from position 0). Stop codons were excluded. The codon-to-amino-acid mapping was derived from Biopython’s Bio.Data.CodonTable.standard_dna_table. RSCU cosine distance between two genomes was calculated as 1 − cos(θ), where cos(θ) is the cosine similarity of their 59-dimensional RSCU vectors (one element per sense codon, excluding methionine, tryptophan, and stop codons). Formally: cosine distance = 1 − (Σ RSCU_a × RSCU_b) / (√Σ RSCU_a² × √Σ RSCU_b²). This was implemented in Python without external libraries and computed directly.

Open reading frames (ORFs) were predicted using NCBI ORFfinder (translation table 1; standard genetic code; https://www.ncbi.nlm.nih.gov/orffinder/; accessed 20 December 2025). Predicted amino acid sequences were analyzed using the NCBI Conserved Domain Database (CDD) via the Conserved Domain Search tool (**Marchler-Bauer and Bryant, 2004**) to identify putative functional domains.

**Table 1.** Quantitative analyses of the assembled viral sequences associated with *Thecaphora thlaspeos*.

| Virus name<br>(abbrev.) | Sequence<br>length<br>(nt) | Mean<br>Reads<br>* | Mean<br>RPKM<br>(± SD)* | Mean<br>Cov.<br>(± SD) | CV<br>(%) | RdRp | | CP | |
| --- | --- | --- | --- | --- | --- | --- | --- | --- | --- |
|  |  |  |  |  |  | nt / aa | CPD<br>family<br>(E-value)<sup>a</sup> | nt /<br>aa | CPD<br>family<br>(E-value)<sup>a</sup> |
| Thecaphora<br>thlaspeos<br>totivirus 1<br>(TtTV1) | 4480 | 2,211 | 14.04<br>± 1.78 | 134.5<br>±<br>38.3× | 12.<br>7 | 2421<br>/<br>806 | pfam0212<br>3 (6.41<br>$\times 10^{-59}$) | 2010<br>/<br>669 | pfam0922<br>0 (1.40<br>$\times 10^{-22}$) |
| Thecaphora<br>thlaspeos<br>totivirus 2<br>(TtTV2) | 4722 | 386 | 2.26 ±<br>0.86 | 22.3 ±<br>11.5× | 37.<br>9 | 2238<br>/<br>745 | pfam0212<br>3 (2.67<br>$\times 10^{-63}$) | 2040<br>/<br>679 | No<br>domains<br>identified |
| Thecaphora<br>thlaspeos<br>totivirus 3<br>(TtTV3) | 4675 | 555 | 3.28 ±<br>1.28 | 32.5 ±<br>18.3× | 38.<br>9 | 2466<br>/<br>821 | pfam0212<br>3 (1.18 ×<br>10<sup>-62</sup>) | 2055<br>/<br>684 | pfam0922<br>0 (1.67 ×<br>10<sup>-10</sup>) |
| Thecaphora<br>thlaspeos<br>totivirus 4<br>(TtTV4) | 4626 | 142 | 0.85 ±<br>0.33 | 8.4 ±<br>4.8× | 39.<br>5 | 2244<br>/<br>747 | pfam0212<br>3 (4.36 ×<br>10<sup>-68</sup>) | 2070<br>/<br>689 | No<br>domains<br>identified |
| Thecaphora<br>thlaspeos<br>totivirus 5<br>(TtTV5) | 4591 | 210 | 1.28 ±<br>0.34 | 12.5 ±<br>5.4× | 26.<br>6 | 2160<br>/<br>719 | pfam0212<br>3 (1.29 ×<br>10<sup>-64</sup>) | 2160<br>/<br>719 | pfam0922<br>0 (2.81 ×<br>10<sup>-6</sup>) |
| Thecaphora<br>thlaspeos<br>eimeriavirus<br>1 (TtEV1) | 5088 | 2,483 | 13.27<br>± 7.34 | 134.3<br>±<br>98.2× | 55.3 | 2478<br>/<br>825 | pfam0212<br>3 (3.61 ×<br>10<sup>-97</sup>) | 2403<br>/<br>800 | pfam0551<br>8 (2.60 ×<br>10<sup>-72</sup>) |
| Thecaphora<br>thlaspeos<br>eimeriavirus<br>2 (TtEV2) | 5296 | 938 | 4.90 ±<br>1.88 | 48.6 ±<br>27.1× | 38.4 | 2673<br>/<br>890 | pfam0212<br>3 (4.27 ×<br>10<sup>-127</sup>) | 2274<br>/<br>757 | pfam0551<br>8 (1.64 ×<br>10<sup>-167</sup>) |
| Thecaphora<br>thlaspeos<br>eimeriavirus<br>3 (TtEV3) | 5080 | 4,115 | 22.80<br>± 7.40 | 222.4<br>±<br>107.1× | 32.4 | 2457<br>/<br>818 | pfam0212<br>3 (9.58 ×<br>10<sup>-124</sup>) | 2277<br>/<br>758 | pfam0551<br>8 (2.50 ×<br>10<sup>-178</sup>) |

<sup>a</sup> Conserved protein domains (CPDs) were identified using the NCBI CD-Search program (database: CDD; E-value cutoff = 0.01). Domain pfam02123 corresponds to RdRps from luteoviruses, totiviruses, and rotaviruses; pfam09220 corresponds to the major capsid protein of the *Saccharomyces cerevisiae* L-A virus; and pfam05518 corresponds to the capsid protein of totiviruses.
<sup>b</sup> BLASTp searches was performed against the nr/pr database (all non-redundant GenBank CDS) using the predicted RdRp amino acid sequences of the putative viruses identified in this study.
\* Summary statistics for each virus across LF2 biological replicates (n = 4). Values are expressed as mean ± SD. CV denotes the coefficient of variation and reflects inter-replicate variability among the eight viruses detected in LF2 libraries.

(Table 1 continued)
| BLASTn search <sup>b</sup> |  |  |  |  |
| --- | --- | --- | --- | --- |
| Best Hit | NCBI GenBank accession | Query cover | Identity | e-value |
| Porphyridium purpureum toti-like virus 2 | CAI5383891.1 | 83 % | 49.85 % | 0.0 |
| Poaceae Liege totivirus 7 | UVG55933.1 | 100 % | 54.55 % | 0.0 |
| Diaporthe helianthi totivirus 2 | WNM95055.1 | 96 % | 51.52 % | 0.0 |
| Umbelopsis<br>gibberispora virus 2 | WGM49131.1 | 100 % | 49.47 % | 0.0 |
| Trichoderma<br>koningiopsis<br>totivirus 1 | QGA70771.1 | 100 % | 53.47 % | 0.0 |
| Mucor hiemalis virus<br>2 | CAE9672664.1 | 99.0 % | 37.40% | 7e-164 |
| Totiviridae sp. | UHS72418.1 | 81 % | 50.14% | 00.0 |
| Totiviridae sp. | UHS72418.1 | 92.0 % | 46.69 % | 00.0 |

### RNA secondary structure and pseudoknot prediction in the overlapping ORF context

The KnotInFrame v2.3.2 algorithm (**Theis et al., 2008**) was used to identify candidate H-type pseudoknots located downstream of canonical −1 programmed ribosomal frameshifting (−1 PRF) slippery heptanucleotides motif (NNN_WWW_H). In parallel, ProbKnot (**Bellaousov and Mathews, 2010**), as implemented in RNAstructure v6.5 (**Reuter and Mathews, 2010**) with a maximum of 10,000 iterations, and IPknot v1.1.0 (**Sato et al., 2011; Sato and Kato, 2022**) were employed as independent algorithms to predict potential pseudoknotted secondary structures. The most probable nested (pseudoknot-free) secondary structures were predicted using the RNAfold algorithm implemented in the ViennaRNA Web Services (**Gruber et al., 2008**), based on the ViennaRNA package v2.5.1 (**Lorenz et al., 2011**), under default thermodynamic parameters (37°C). Analyses were performed via the Vienna RNA Web Server (http://rna.tbi.univie.ac.at/; accessed 25 February 2026). For each predicted structure, folding free energies (ΔG) of predefined RNA secondary structures were calculated using the Energy Function 2 (efn2) program (RNAstructure v6.5; **Reuter and Mathews, 2010**), which applies nearest-neighbor thermodynamic parameters. To avoid potential bias arising from differences in sequence length, ΔG values were normalized by sequence length (kcal/mol/nt) prior to comparative analyses (**Chan and Ding, 2008; Jacquat et al., 2024**). To further assess relative thermodynamic stability between alternative conformations, a dimensionless Δrel metric was calculated as Δrel = (ΔG_nested − ΔG_pk) / |ΔG_nested|, where ΔG_pk corresponds to the predicted pseudoknotted structure and ΔG_nested to the most stable alternative nested conformation. Positive Δrel values indicate a thermodynamic advantage of the pseudoknotted structure over the nested alternative.

### RdRp dataset retrieval and curation for phylogenetic analysis

RNA-dependent RNA polymerase (RdRp)–like amino acid sequences were retrieved from the NCBI RefSeq Protein database using BLASTp with an E-value cutoff of 1 × 10⁻⁵. The RdRp sequence of the type species *Totivirus ichi* (representative member: Saccharomyces cerevisiae virus L-A; GenBank accession AAA50508) was used as the query. A total of 49 sequences were recovered and combined with 32 RdRp sequences corresponding to ICTV-recognized Totivirus species (**Supplementary Table S1**). To reduce possible redundancy, sequences were clustered using CD-HIT v4.8.1 (**Li and Godzik, 2006**) at a 90% pairwise amino acid identity threshold. Sequences shorter than 600 amino acids were excluded as putatively truncated. For RefSeq entries annotated as capsid protein (CP)–RdRp fusion proteins, the N-terminal CP region was manually trimmed based on alignment with the reference S. cerevisiae virus L-A genome. The resulting curated dataset comprised 56 nonredundant RdRp sequences (**Supplementary Data S1**).

An equivalent procedure was applied to retrieve RdRp sequences related to the genus *Eimeriavirus*. The RdRp sequence of the type species *Eimeriavirus ichi* (Eimeria tenella RNA virus 1; GenBank accession AIW58883) was used as the query, yielding 73 sequences, including three ICTV-recognized members (**Supplementary Table S1**). After clustering at 90% identity, 66 nonredundant sequences were retained (**Supplementary Data S2**).

To eliminate redundancy between the *Totivirus* and *Eimeriavirus* datasets prior to multiple sequence alignment and phylogenetic inference, both datasets were pooled and subjected to an additional CD-HIT clustering step at 90% identity. This procedure removed 19 and 6 sequences from the *Totivirus* and *Eimeriavirus* subsets, respectively. Overlap between the two BLASTp-derived datasets was expected due to sequence similarity among evolutionarily related genera. Two ICTV-recognized RdRp sequences from the genus *Megatotivirus* (**Supplementary Table S1**) were included as outgroup taxa for phylogenetic reconstruction. Members of this genus belong to the suborder *Betatotivirineae*, which is phylogenetically distinct from *Alphatotivirineae*, the suborder comprising *Totivirus* and *Eimeriavirus*. The final dataset consisted of the eight virus-like RdRp sequences identified in this study, 97 curated reference sequences, and two outgroup sequences (**Supplementary Data S3**).

### Phylogenetic reconstruction

The final dataset used for RdRp phylogenetic inference (**Supplementary Data S3**) exhibited a global all-versus-all mean pairwise amino acid identity of 29.5% (SD = 6.2%). Pairwise amino acid sequence identities were calculated using BLASTp v2.12.0+ (**Camacho et al., 2009**). Local all-versus-all comparisons were performed after removal of self-hits and redundant reciprocal matches. At this level of divergence, RdRp sequences from ssRNA viruses are still expected to retain sufficient phylogenetic signal to resolve relationships at the family and genus levels when appropriate alignment strategies are employed. Multiple sequence alignment was performed using MAFFT v7.511 (**Katoh et al., 2002; Katoh et al., 2019**) under the FFT-NS-i iterative refinement algorithm in combination with the Database of Aligned Structural Homologs (MAFFT-DASH). This strategy was selected as suitable for the observed level of sequence similarity. The MAFFT-DASH alignment comprised 107 sequences and 2,286 alignment columns. Of these, 16.2% were constant sites, 26.8% were singleton sites, 57.0% were parsimony-informative sites, and 92.9% represented distinct site patterns.

Maximum likelihood (ML) phylogenetic inference was conducted by IQ-TREE version 2.0.7 (**Minh *et al*., 2020**). The best-fitting evolutionary matrix was automatically selected with ModelFinder (**Kalyaanamoorthy *et al*., 2017**) based on standard information-theoretic criteria as implemented in IQ-TREE 2. Node statistical support was assessed using the ultrafast bootstrap approximation (UFBoot2; **Hoang *et al*., 2018**) based on 1,000 replicates. The inference was performed at IQ-TREE web server (**Trifinopoulos et al., 2016**). The best-fitting substitution model was LG+F+R9. Estimated site-proportion and rate parameters for the FreeRate model are: (0.018, 0.008) (0.021, 0.053) (0.027, 0.141) (0.071, 0.265) (0.103, 0.416) (0.164, 0.646) (0.129, 0.890) (0.250, 1.2075) (0.217, 1.894). Total tree length (sum of branch lengths): 104.44

## Results

### Eight bona fide virus-like sequences associated with Thecaphora thlaspeos were identified in transcriptomic libraries

The re-analysis of the sequencing libraries, based on filtering, RNA assembly, bulk BLASTX against NCBI refseq virus proteins and transcript curation produced 13 assembled virus-like RNA sequences that were detected almost exclusively in *T. thlaspeos* libraries (**Figure 1; supplementary table S2**), and based on multiple analyses described below, were assigned to Thecaphora thlaspeos totivirus 1–5 (TtTV1–5) and Thecaphora thlaspeos eimeriavirus 1–3 (TtEV1–3). In strain LF2, virus-mapped reads per library ranged from 7,463 to 20,277, representing 0.0221% to 0.0481% of the total output. The LF1 library contained 3,165 virus reads, accounting for 0.0245% of 12.9 million total reads. In contrast, plant control libraries yielded only 5 to 34 virus-associated reads each (0.0000% to 0.0001%), consistent with low-level index hopping or barcode cross-contamination rather than genuine infection. Mapping analysis of DNA sequencing reads from the same bioproject and samples against these putative viral sequences provided additional evidence for their RNA nature. Across all eight DNA libraries totaling approximately 45 million paired-end reads, zero reads mapped to any of the eight viral assemblies. This complete absence of viral sequences in DNA preparations confirms that the identified viruses represent authentic RNA viruses rather than endogenous viral elements, retrotransposons, or DNA contaminants. The comprehensive nature of this negative result, spanning multiple independent DNA extractions and sequencing runs, provides robust evidence against chromosomal integration of these viral sequences

**Figure 1.**
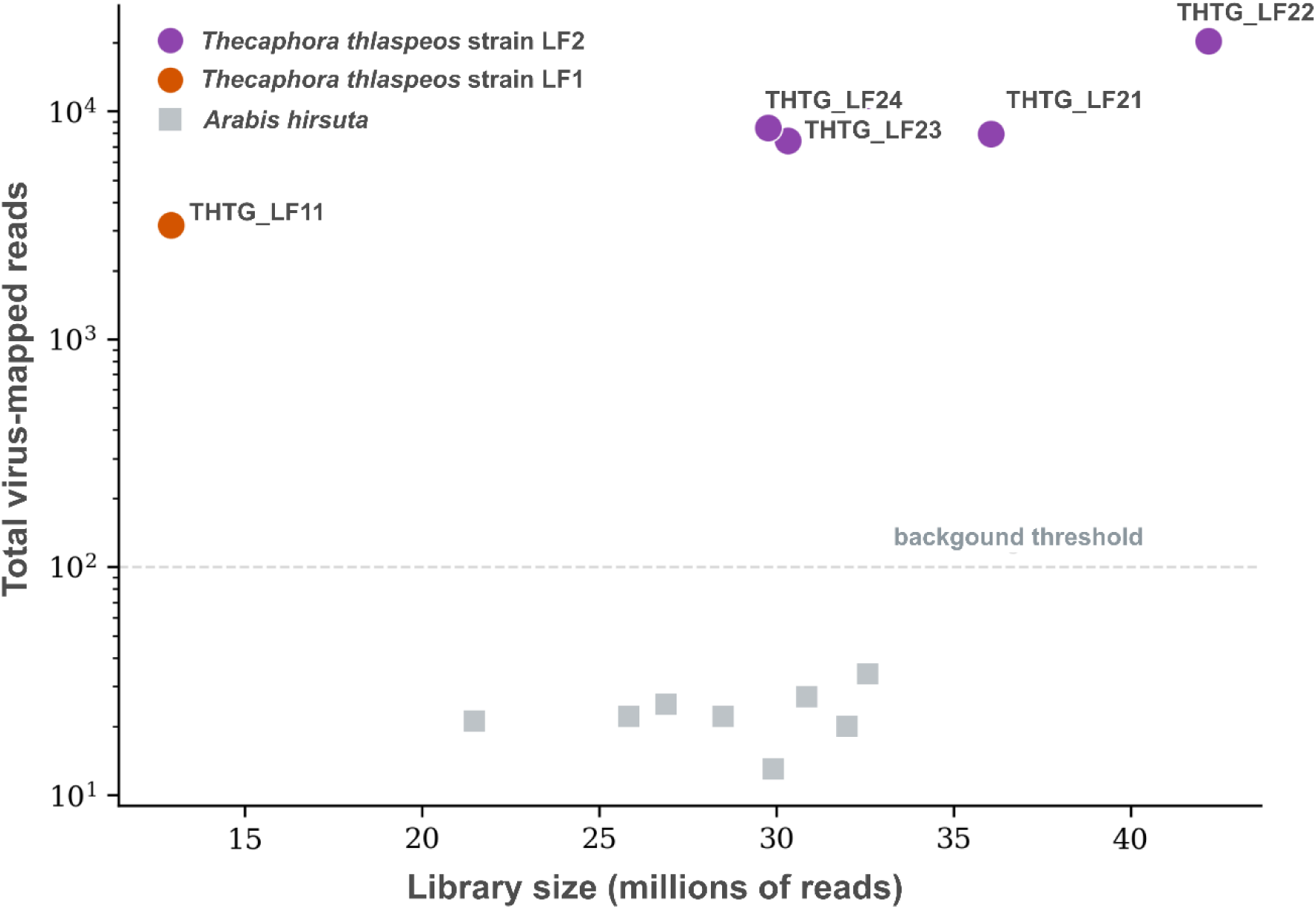
Virus enrichment in fungal vs plant libraries. Scatter plot of total virus-mapped reads (Y-axis, log scale) as a function of library size in millions of reads (X-axis). Circles represent fungal libraries colored by strain (purple = LF2, orange = LF1); squares represent *A. hirsuta* plant controls (grey). All fungal libraries cluster at 10³–10⁴ virus reads, while plant controls fall at 10⁰–10¹, consistent with background-level index-hopping. The dashed line at 100 reads indicates an approximate threshold separating genuine viral presence from sequencing spillover. The heatmap of genome coverage of the figure 3 further illustrates the distribution of viral reads across libraries (Figure 3).

Pairwise nucleotide identity comparisons among all 13 viral sequences detected (tentatively assigned to eight virus species across two fungal strains) revealed a clear separation between intra- and inter-species divergence levels (**Figure 2**). Within-species strain pairs (LF1 vs. LF2) exhibited identities ranging from 87.74% (TtTV3) to 93.68% (TtTV5). By contrast, the eight assembled sequences yielded identities between 34% and 49%. A block structure was evident in the identity matrix, with the three eimariaviruses (TtEV1–TtEV3) forming one cluster (within-group identities 40–48%) and the five totiviruses (TtTV1–TtTV5) forming another (39–49%), while between-family comparisons were uniformly lower (34–40%). Five of the eight viruses, TtEV1, TtTV1, TtTV3, TtTV4, and TtTV5, were detected in both fungal strains, whereas TtEV2, TtEV3, and TtTV2 were recovered only from LF2 libraries; however, because LF1 is represented by a single sequencing library, this absence should not be interpreted as confirmed virus loss without additional sampling.

**Figure 2.**
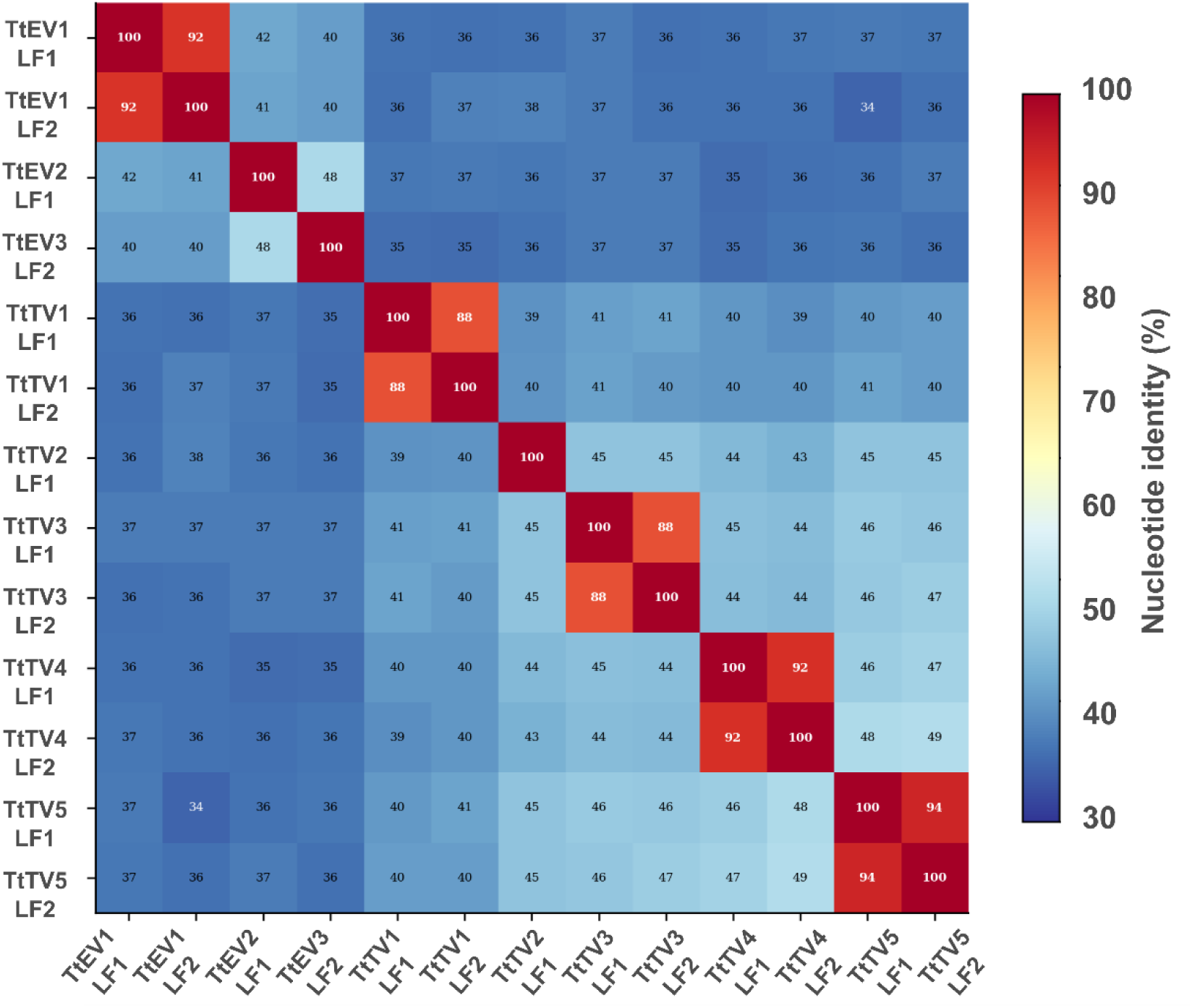
Pairwise nucleotide identity matrix of *T. thlaspeos* RNA virus genomes. Heatmap displaying pairwise nucleotide identities (%) among all 13 viral genome sequences, calculated by direct positional comparison of ungapped sequences with identity expressed as the proportion of matching positions over the length of the longer sequence. Diagonal values (100%) represent self-comparisons.

The deduced RdRp amino-acid sequences of these eight assembled virus-like RNA sequences revealed presence of conserved motifs characteristic of the viral RdRp superfamily (Pfam: pfam02123), which includes polymerases from members of the families *Totiviridae*, *Reoviridae*, and *Luteoviridae*. Additionally, conserved domains corresponding to totivirus capsid proteins were detected (Pfam: pfam05518 and pfam09220) (**Table 1**). However, a clear distinction was evident between two groups based on GC content, sequence length and genomic architecture. Specifically, five sequences (mean GC content 53.5%) ranged from 4,480 to 4,722 nt and contained two partially overlapped ORFs. In contrast, three sequences with higher GC content (mean 56.8%) ranged from 5,080 to 5,296 nt and harbored two ORFs separated by a conserved AUGA tetranucleotide motif, typical of a stop–restart translational mechanism. These features are consistent with the canonical genome organization of *Totiviruses* and *Eimariaviruses*, respectively. Further details of this genomic architecture are presented in the section “Structural characterization of viruses associated with Thecaphora thlaspeos”.

Phylogenetic inference based on RdRp sequences constitutes the primary ICTV criterion for classification within the order *Ghabrivirales* (**ICTV, 2024).** To taxonomically position the eight virus-like RNA sequences, a phylogenetic reconstruction was performed following the ICTV framework (see Materials and Methods). Five of these sequences clustered with maximal statistical support (100% UFboot) with members of the genus *Totivirus* and closely related sequences. Similarly, the other three sequences formed a highly supported clade (100% UFboot) with members of the genus *Eimeriavirus* and their close relatives (**Figure 3**). Based on this clustering, the identified virus-like RNA sequences were provisionally classified as derived from viral genomes provisionally designated as Thecaphora thlaspeos totivirus 1–5 (TtTV1–5) and Thecaphora thlaspeos eimeriavirus 1–3 (TtEV1–3) (**Table 1**). Furthermore, amino acid identities (BLASTp) of the deduced RdRp ranged from 37.40% to 54.55% (coverage 80% - 100%) with the closest known viruses (**Table 1**). These values are well below the 70% threshold used by ICTV for species demarcation in this group (**ICTV, 2024**), supporting the classification of these viruses as member of putative novel species.

**Figure 3.**
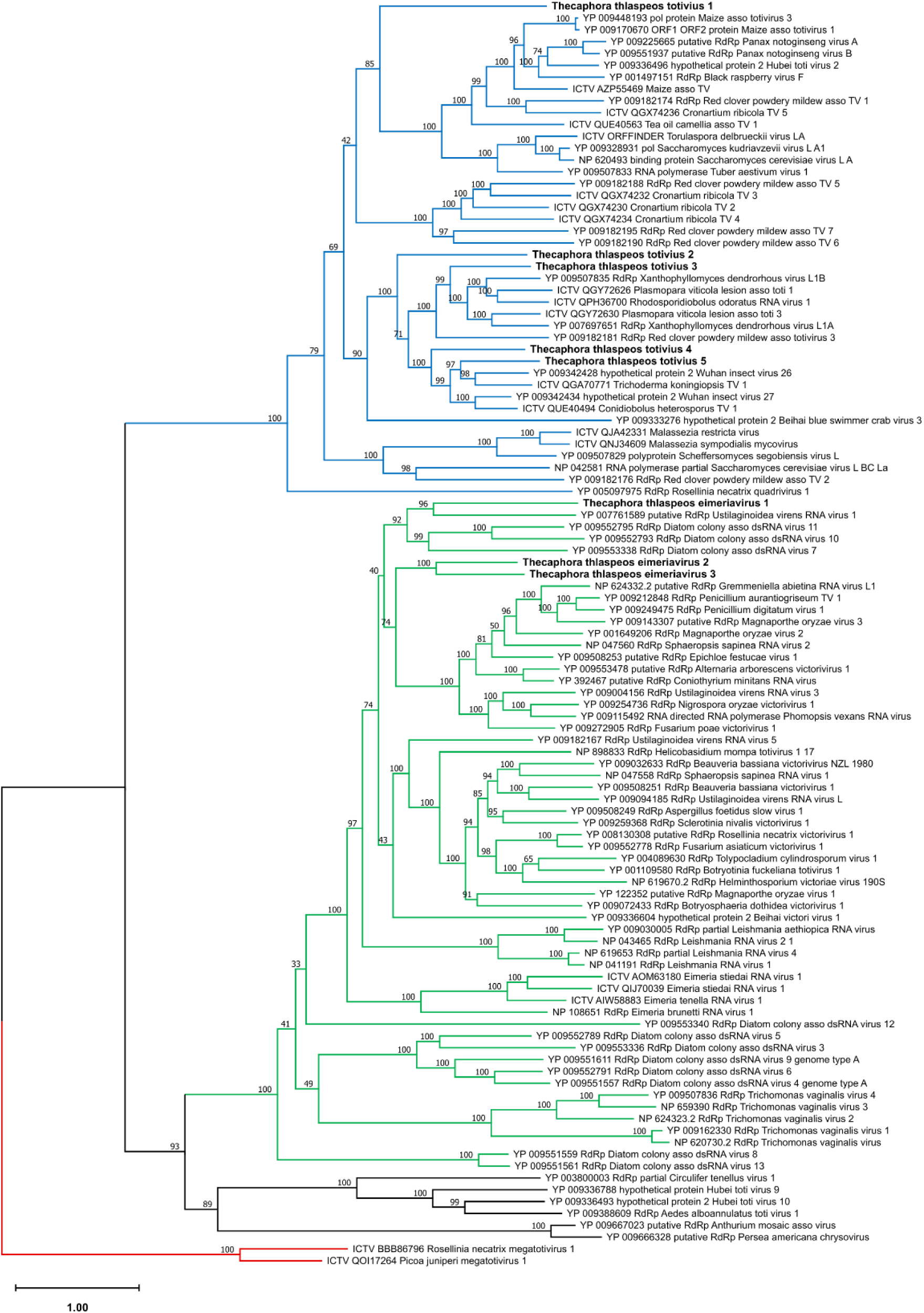
Maximum-likelihood phylogeny of RdRp amino-acid sequences. The tree was inferred from a MAFFT-DASH alignment under the LG+F+R9 substitution model, as selected according to model-fitting criteria (see Materials and Methods). Branch support values correspond to ultrafast bootstrap percentages based on 1,000 replicates. Terminal labels indicate NCBI accession numbers followed by virus names; ICTV-recognized species are designated as “ICTV”, and viruses identified in this study are shown in bold. Two well-supported monophyletic lineages corresponding to the family *Orthototiviridae* (blue branches) and *Pseudototiviridae* (green branches) are recovered. Members of the *Megatotiviridae* family (red branches) were used as outgroup to root the tree. The scale bar represents the median number of amino-acid substitutions per site.

### Viral abundance and community composition

In strain LF2, viruses belonging to the genus *Eimariavirus* (TtEV1–TtEV3) collectively accounted for 56–61% of total virus-mapped reads across the four biological replicates, whereas members of the genus *Totivirus* (TtTV1–TtTV5) comprised the remaining 39–44%. In LF1, the overall genus-level distribution appeared similar, with *Eimariavirus* representing approximately 59% of virus reads. However, this proportion was driven entirely by TtEV1, as TtEV2 and TtEV3 were completely absent from this strain (**Supplementary Table S2**).

TtEV3 from the fungal strain LF2 exhibited the highest viral abundance with a mean RPKM of 22.80 ± 7.40 and a mean genome coverage of 222.4 ± 107.1× (**Table 1**, **Figure 4**). TtTV1 was the second most abundant virus, showing a mean RPKM of 14.04 ± 1.78 and the lowest inter-replicate variability among all eight viruses (CV = 12.7%), indicating stable accumulation across biological replicates (**Table 1**, **Figure 4**). TtEV1 displayed a mean RPKM of 13.27 ± 7.34 but substantially higher variability (CV = 55.3%), largely driven by elevated read counts in replicate THTG_LF22, which also had the largest library size and the highest number of virus-mapped reads. The remaining viruses (TtEV2 and TtTV2–TtTV5) were detected at lower abundance, with mean RPKM values ranging from 0.85 to 4.90 (**Table 1**, **Figure 4**).

**Figure 4.**
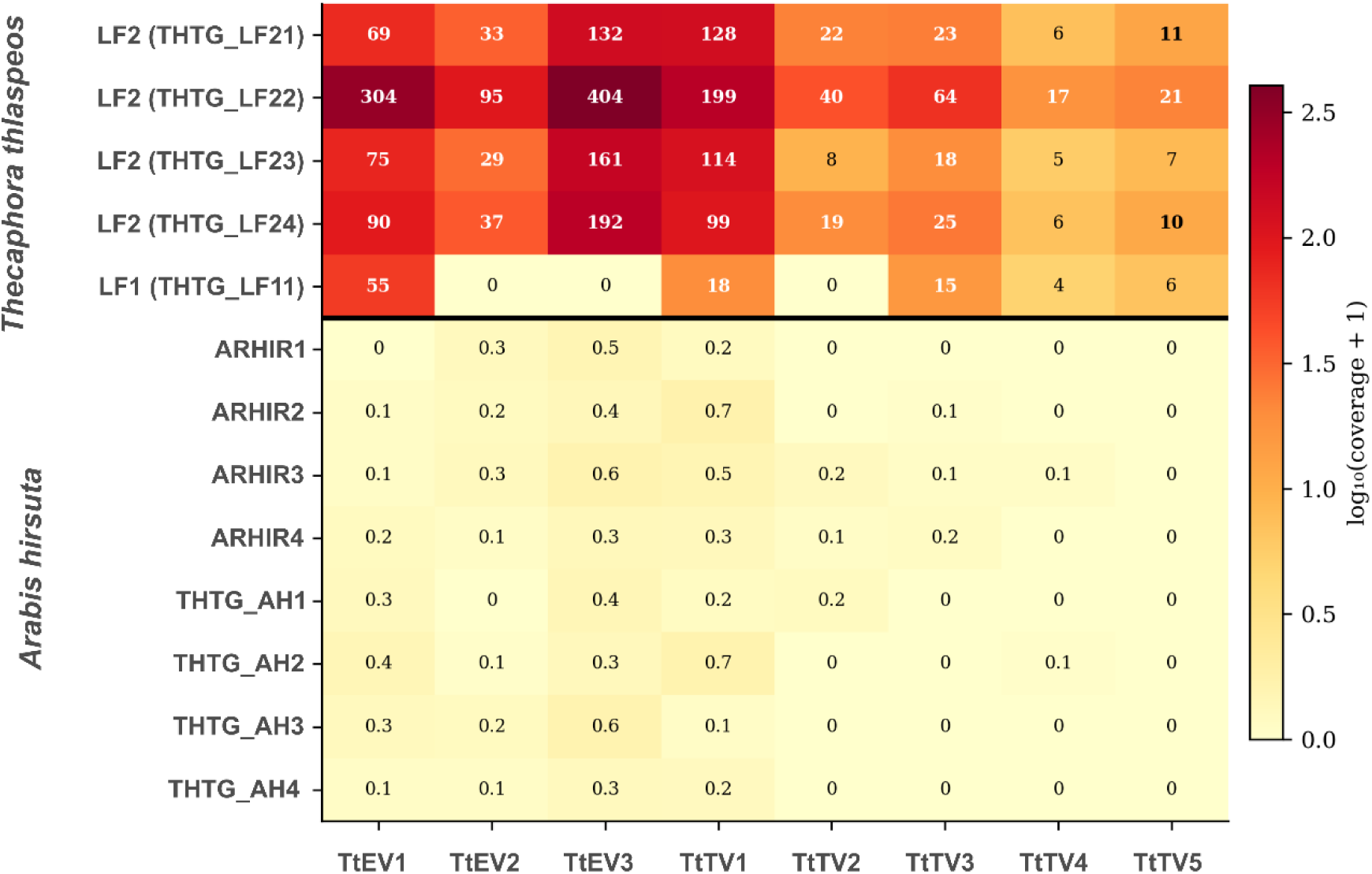
Heatmap of mean genome coverage across all libraries. Heatmap displaying log₁₀(coverage + 1)-scaled mean genome coverage for each virus (columns) across all 13 RNA-seq libraries (rows). The upper block shows the five fungal mycelium libraries; the lower block shows the eight *A. hirsuta* plant controls. Numeric values indicate coverage depth (×).

The viral profile of fungal strain LF1 differed markedly from that of strain LF2. As only five of the eight viruses were detected: TtEV1 (RPKM = 28.29, the highest single-sample value observed for any virus in the dataset), TtTV1 (9.48), TtTV3 (7.69), TtTV4 (2.03), and TtTV5 (2.95). In the library with the highest sequencing depth (THTG_LF22), genome coverage reached 404.22× for TtEV3 and 198.54× for TtTV1 (**Figure 4; Supplementary Table S2**). Coverage in LF1 was lower (3.9–55.1×), reflecting both the shorter single-end read length (151 bp versus ∼270 bp) and the smaller library size, yet remained sufficient to support the presence of the viruses detected in this dataset. Consistent with adaptation to the host translational environment, cosine distances between viral and host relative synonymous codon usage (RSCU) profiles were extremely small (0.0084–0.0109), indicating highly similar codon preferences. Indicating an almost complete overlap in codon usage preferences. Such a high degree of similarity strongly suggests translational optimization of viral genomes to the host machinery, consistent with long-term virus–host coadaptation.

### Genome-wide divergence patterns between strains and signatures of purifying selection

Sliding window analysis of nucleotide divergence revealed a non-uniform distribution of sequence differences across viral genomes (**Figure 5A**). Genome-wide mean divergence ranged from 6.4% to 12.4%, consistent with overall pairwise identity estimates (**Figure 2**). Specifically, the five viruses present in both LF1 and LF2 strains exhibit moderate levels of nucleotide divergence, with pairwise identities ranging from 87.74% (TtTV3) to 93.68% (TtTV5). The eimariavirus TtEV1 showed 91.51% identity between strains, positioning it in the middle of the observed range and indicating similar evolutionary pressures across viral families. Across all viruses, conserved regions were interspersed with localized peaks of higher divergence, indicating that sequence variation is structured rather than evenly distributed. In most cases, the RdRp-encoding region exhibited greater variability than the CP (**Figure 5A**), suggesting differential evolutionary constraints across genomic regions. No extended regions of extreme divergence were observed. Mutation density analysis revealed substantial variation in the number of accumulated differences, ranging from 290 single nucleotide polymorphisms (SNPs) in TtTV5 to 573 SNPs in TtTV3. When normalized by genome length, SNP densities varied from 63.2 to 122.6 substitutions per kilobase, suggesting heterogeneous evolutionary rates across the viral community (**Figure 5B**).

**Figure 5.**
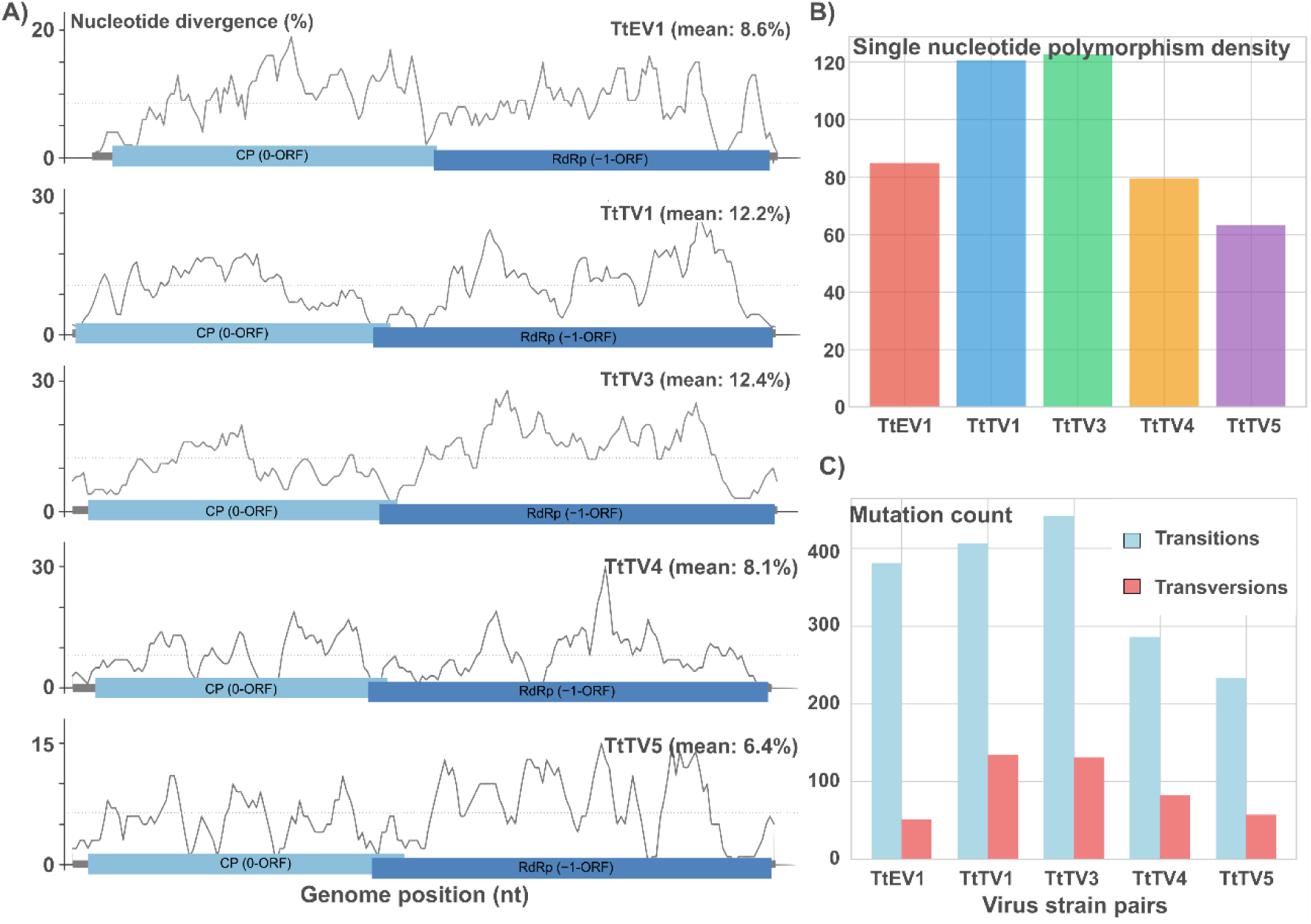
Mutation landscape and divergence profiles of the five shared *T. thlaspeos* viruses between strains LF1 and LF2. (**A**) Single nucleotide polymorphism (SNP) density, expressed as SNPs per kilobase, for each virus strain pair. Blue and dark blue boxes represent CP (0-ORF) and RdRp (−1-ORF), respectively. (**B**) Genome-wide nucleotide divergence between LF1 and LF2 viral sequences. Sliding window analysis of nucleotide divergence was performed for each virus using pairwise alignments generated with MAFFT v7.511. Divergence was calculated as the proportion of mismatched nucleotide positions between LF1 and LF2 sequences within 100 bp windows, with a step size of 25 bp. Values are plotted across relative genome positions. Dashed horizontal lines indicate the mean genome-wide divergence for each virus. Genomic organization is shown above each plot, with the capsid protein (CP; 0-ORF) and RNA-dependent RNA polymerase (RdRp; −1 ORF) regions indicated. (**C**) Absolute counts of transitions (light blue) and transversions (red) for each virus strain pair. All five viruses exhibit transition-biased mutation spectra, consistent with the expected mutational bias of RNA-dependent RNA polymerases.

Despite moderate nucleotide divergence, amino acid sequences remain highly conserved across all viral proteins. Coat protein identities ranged from 98.75% to 99.85%, while RdRp sequences showed 97.32% to 99.44% identity. The fusion proteins generated through ribosomal frameshifting maintained 98.22% to 99.47% identity between strains. This pattern is accompanied by a marked excess of transitions over transversions (**Figure 5C**), which is consistent with the accumulation of synonymous substitutions. Transition to transversion ratios ranged from 3.03 to 7.47, with the eimariavirus TtEV1 showing the highest transition bias. These values exceed the neutral expectation of approximately 2.0.

This pattern—moderate nucleotide divergence, strong amino acid conservation, and near-identical codon usage relative to the host—supports a model in which synonymous mutations accumulate while maintaining both protein function and translational efficiency, consistent with long-term purifying selection acting at multiple levels.

### Genomic architecture of totiviruses and eimeriaviruses associated to Thecaphora thlaspeos

A detailed nucleotide sequence analysis indicates that the putative genomes of TtTV1–5 conform to the canonical *Totivirus* genomic architecture (**Figure 6 A-E**), comprising two cis-contiguous, partially overlapping 5′→3′ open reading frames (ORF1 and ORF2) and a putative −1 programmed ribosomal frameshifting (−1 PRF) mechanism enabling translation of a CP–RdRp fusion protein (Gag–Pol polyprotein). In the L-A totivirus of Saccharomyces cerevisiae, experimental evidence demonstrated that efficient −1 ribosomal frameshifting requires typically a pseudoknot (stimulatory RNA structure) located immediately downstream of the slippery heptanucleotide motif XXX_YYY_Z (**Dinman et al., 1991**); more generally represented in IUPAC notation as NNN_WWW_H, in which N denotes any nucleotide, W represents A or T, and H corresponds to any nucleotide except G. Firstly, a multiple sequence alignment (MSA) including 32 ICTV-recognized *Totivirus* exemplars revealed a highly conserved heptanucleotide motif located near the 3′ terminus of ORF1 (**Figure 7A**). In this study, Saccharomyces cerevisiae virus L-A, in which −1PRF is mediated by the slippery motif GGGTTTA (**Dinman et al., 1991**), was used as a reference. The multiple sequence alignment revealed that the predominant conserved heptamer among the analyzed sequences corresponds to GGGTTTT, with GG(A/G)TTTT representing the most frequent variant (**Figure 7B**).

**Figure 6.**
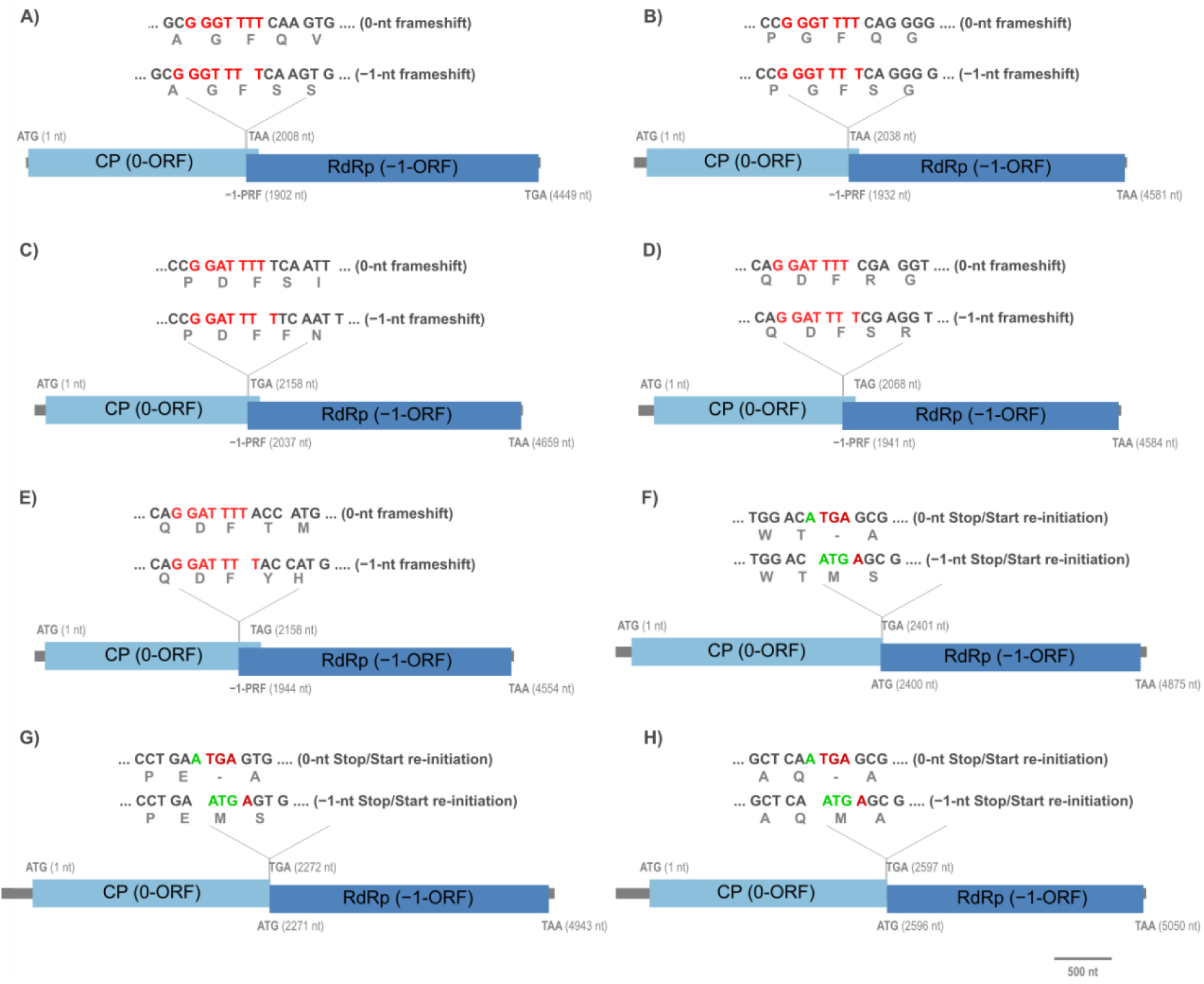
Genomic organization and translational strategies of Thecaphora thlaspeos-associated viruses. (A–E) Thecaphora thlaspeos associated totiviruses 1–5 utilize −1 programmed ribosomal frameshifting (−1 PRF) to express the RNA-dependent RNA polymerase (RdRp). Conserved heptanucleotide slippery sequences (red) promote ribosomal shifting from the 0-frame (top) to the −1-frame (bottom) upstream of the CP stop codon, resulting in CP–RdRp fusion proteins. (F–H) Thecaphora thlaspeos associated eimeriaviruses 1–3 employ a termination–reinitiation (Stop/Start) mechanism. These genomes contain a characteristic tetranucleotide overlap (ATGA), where the shared adenine (green) constitutes both the third nucleotide of the penultimate CP codon and the first nucleotide of the RdRp start codon. Translation of RdRp occurs following CP termination at the adjacent stop codon (red). Blue and dark blue boxes represent CP (0-ORF) and RdRp (−1-ORF), respectively. Amino acids corresponding to the 0-frame and −1-frame translations are shown above and below the nucleotide sequences. Coordinates (in parentheses) denote nucleotide positions relative to the CP start codon (nt 1). Scale bar = 500 nt.

**Figure 7.**
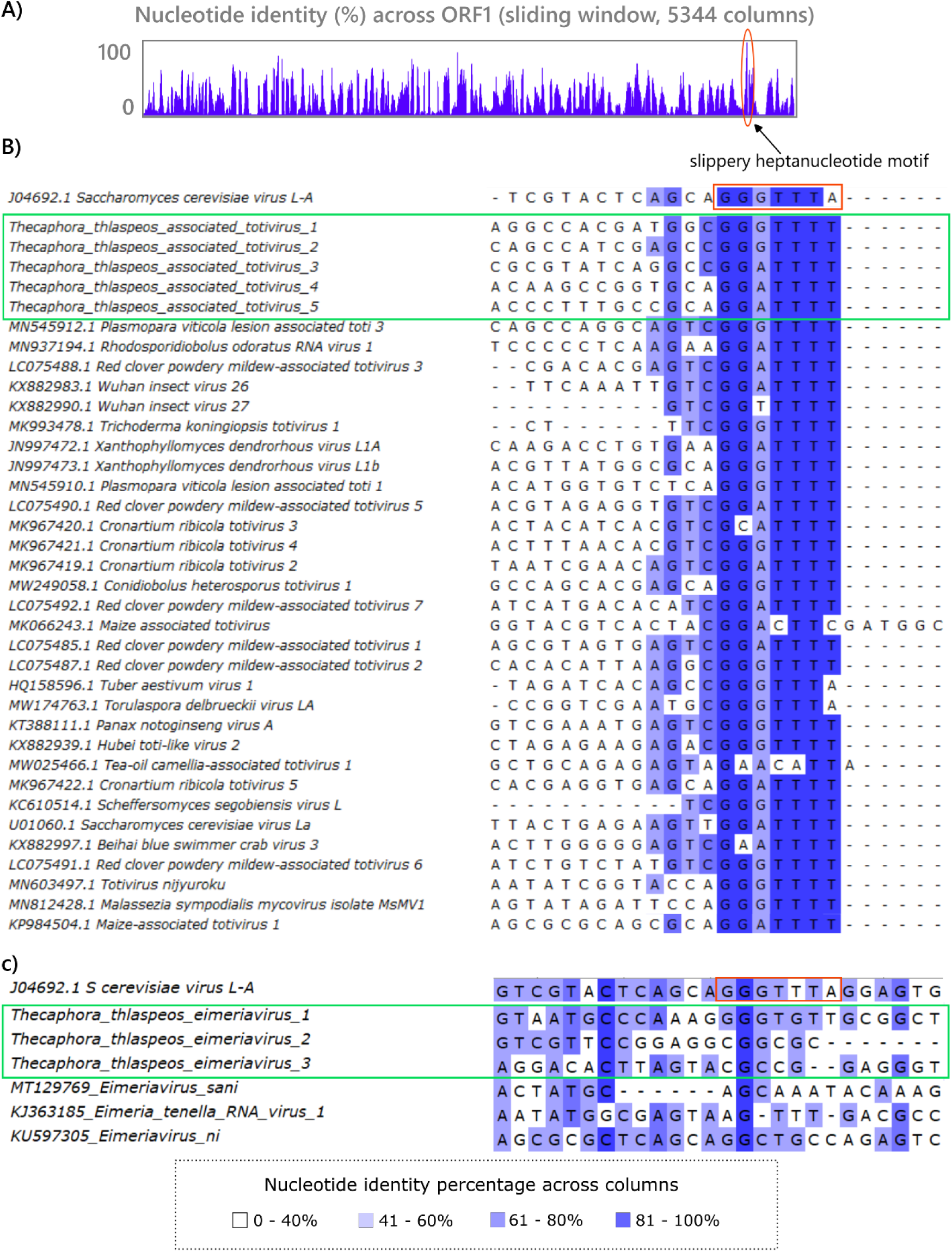
Conservation and sequence context of the putative −1 programmed ribosomal frameshifting (−1 PRF) slippery site in *Thecaphora thlaspeos*–associated viruses. **(A)** Sliding-window analysis of nucleotide identity across ORF1 (5,344 alignment columns) reveals a marked conservation peak near the 3′ terminus corresponding to the predicted slippery heptanucleotide motif. **(B)** Multiple sequence alignment of representative *Totivirus* sequences, including TtATV1–5 (green box) and ICTV-recognized species, showing strong conservation of the canonical XXXYYYZ slippery motif (boxed in red). Nucleotide identity across columns is indicated by a blue gradient. **(C)** Alignment of representative *Eimeriavirus* sequences, including *Thecaphora thlaspeos*–associated eimeriaviruses 1–3 (green box). The genome of *Saccharomyces cerevisiae* virus L-A was included as a reference to indicate the position of the canonical slippery heptamer; however, the associated eimeriaviruses do not exhibit conservation of this region, indicating the absence of a preserved totivirus-like slippery motif. The color scale denotes percentage nucleotide identity per column.

To investigate the potential presence of −1 PRF stimulatory elements, candidate pseudoknotted RNA secondary structures were analyzed downstream of conserved slippery heptanucleotide motifs. Multiple computational approaches were used to predict both knotted and nested conformations, and their thermodynamic stability was comparatively evaluated. Detailed descriptions of the algorithms, energy calculations, normalization procedures, and the Δrel metric are provided in the Materials and Methods section. Pseudoknot prediction was dependent on both the algorithm applied and the length of the sequence window analyzed, highlighting the structural plasticity of this genomic region. Accordingly, the reported results represent a systematic exploration of alternative conformations using complementary computational strategies.

Prediction of RNA secondary structures downstream of the slippery motifs identified candidate pseudoknotted conformations in all five totiviruses (**Figure 8; Supplementary_data_S5 – S9**). For each genome, two candidate pseudoknots were detected within the expected 3′ region relative to the frameshift site. The predicted topologies and stability estimates varied depending on the algorithm and sequence window analyzed. In all cases, the pseudoknotted conformations were thermodynamically less stable than the corresponding nested structures, as reflected by negative Δrel values. However, the predicted free energy differences were moderate, indicating that pseudoknotted conformations remain energetically accessible within the structural ensemble.

**Figure 8.**
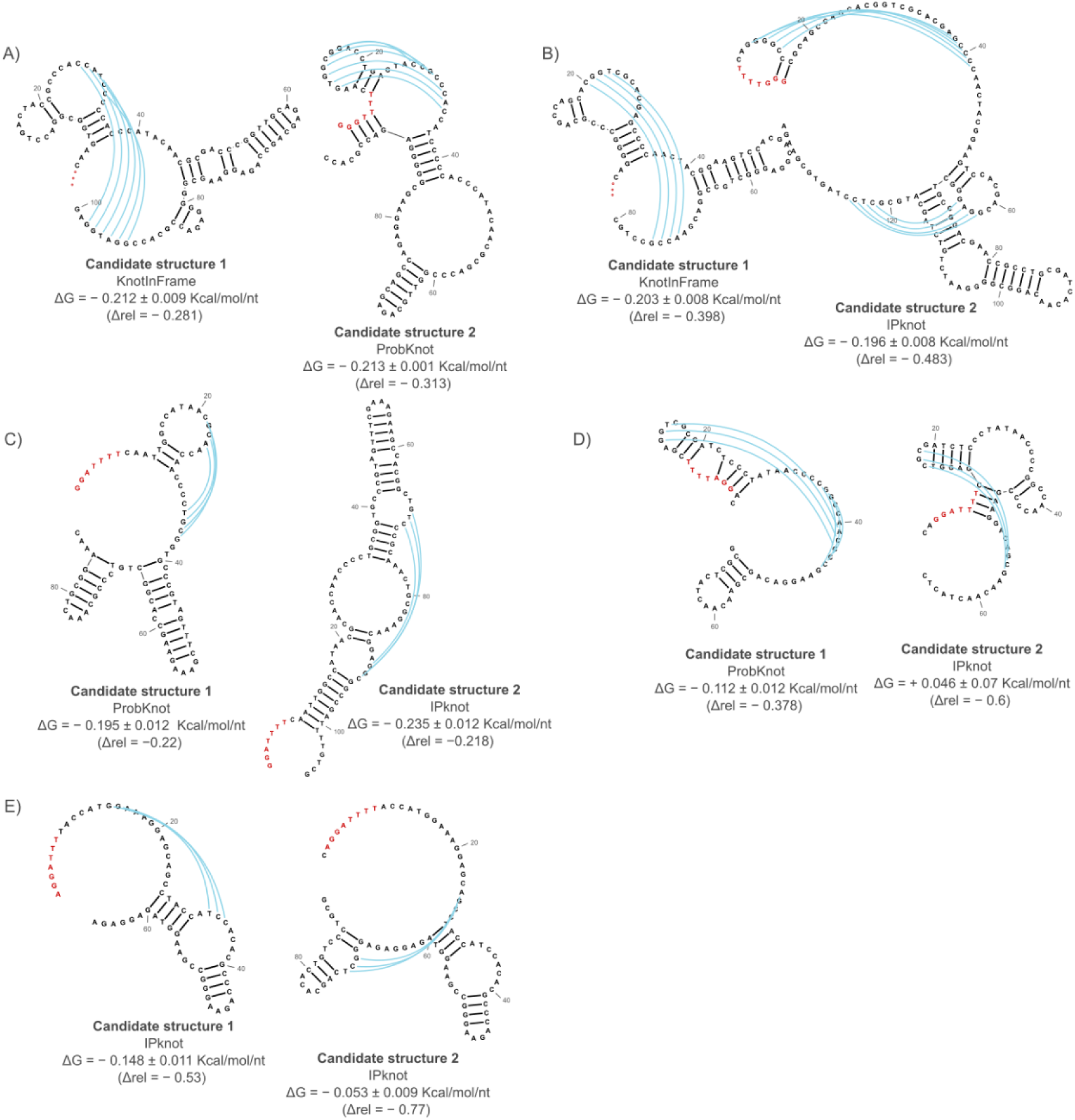
Predicted downstream RNA secondary structures associated with −1 programmed ribosomal frameshifting (−1 PRF) in *Thecaphora thlaspeos*-associated totiviruses 1–5. (A–E) Candidate pseudoknotted RNA structures identified downstream of the slippery heptanucleotide motifs. For each virus, the most thermodynamically favorable structures predicted by KnotInFrame, ProbKnot, and/or IPknot are shown. Blue arcs indicate base-pairing interactions involved in pseudoknot formation. Slippery sequences are highlighted in red. For each candidate structure, normalized minimum free energy (ΔG; kcal/mol/nt) and the relative stability metric (Δrel) are indicated. Δrel was calculated as (ΔG_nested − ΔG_pk) / |ΔG_nested|, where ΔG_pk corresponds to the pseudoknotted conformation and ΔG_nested to the most stable nested (pseudoknot-free) structure. Negative Δrel values indicate reduced thermodynamic favorability of the pseudoknotted structure relative to the alternative nested conformation.

The *Thecaphora thlaspeos* eimeriaviruses 1–3 (**Figures 6F–H**) showed no detectable conservation of a canonical slippery heptanucleotide motif in the region corresponding to the −1 PRF site described for totiviruses (**Figure 7C**). Instead, their genomes comprise two partially overlapping ORFs separated by a highly conserved tetranucleotide ATGA junction. This motif is consistent with a canonical stop/start termination–reinitiation signal, in which the shared adenine functions both as the third nucleotide of the upstream CP ORF stop codon and as the first nucleotide of the downstream RdRp initiation codon (ATG). This genomic arrangement is consistent with a translational coupling mechanism, ensuring that RdRp expression depends on termination of the upstream ORF.

Given the phylogenetic proximity of these viruses to Helminthosporium victoriae virus 190S, in which stop/restart translational recoding has been experimentally demonstrated (**Li et al., 2011**), we used this system as a reference to explore potential structural determinants associated with reinitiation. However, no similar pseudoknot structures or conserved RNA structural patterns were predicted in the region surrounding the ORF1 termination site using ProbKnot and IPknot. The termination–reinitiation in these viruses may depend on alternative, possibly minimal or transient sequence elements that remain unresolved by current in silico approaches.

While these observations are consistent with distinct translational recoding strategies in totiviruses and eimeriaviruses, experimental validation would be required to confirm the underlying molecular mechanisms, which is beyond the scope of the present study

## Discussion

### Virus infection evidence and quantitative aspects

In this study, we report the first viruses in the biotrophic *brassicaceae*-specialist smut fungus *T. thlaspeos*, substantially expanding the known mycovirome of smut fungi in the *Ustilaginomycotina*, a subphylum that remains comparatively underexplored from a virological perspective. Evidence for mycovirus infection derives from transcriptomic data generated from fungal mycelium and from the host plant used as a control. The relative abundance of reads mapped to viral genomes, sequencing depth, genome coverage, and patterns of RSCU are consistent with active viral replication, although direct experimental evidence remains to be obtained.

The absence of viral sequences in DNA libraries provides relevant evidence of the RNA nature of the detected viruses excluding the possibility of endogenous viral elements or viral like retrotransposon contamination. This distinction is particularly important in fungal systems where endogenous viral-like sequences can complicate virus discovery efforts (**Kondo et al., 2013**). The comprehensive DNA mapping approach employed here represents a best practice for confirming the authenticity of RNA virus discoveries in genomic datasets, as endogenous viral elements would be detectable in both RNA and DNA preparations (**Feschotte and Gilbert, 2012**). This validation strengthens confidence in the biological relevance of the identified viral sequences and their classification as active RNA viruses within the *T. thlaspeos* virome.

Phylogenetic placement based on RdRp sequences strongly supports classifying these viruses within the genus *Eimeriavirus* (*Pseudototiviridae*) and *Totivirus* (*Orthototiviridae*). Although totiviruses have been reported in the corn smut *Ustilago maydis* (**Wickner et al., 2013**) and diverse mycoviruses have been described in the rice false smut pathogen *U. virens* (**Jiang et al., 2015; Zhong et al., 2017**), the presence of a complex multi-virus community comprising both *Eimariavirus* and *Totivirus* elements in a *brassicaceae-*infecting smut fungus has not, to our knowledge, been reported.

The proportion of virus-mapped reads in fungal-derived libraries ranged from 0.0221% to 0.0481% of total reads, whereas plant-derived libraries contained only 0.0000–0.0001%. This approximately three-order-of-magnitude disparity in viral read counts between fungal and plant libraries is inconsistent with genuine viral infection of plant tissues. Instead, the minimal counts detected in plant controls are best explained by background index hopping or barcode cross-contamination. Index hopping is a well-characterized artifact of Illumina sequencing platforms, particularly those employing patterned flow cells and exclusion amplification, where read misassignment rates typically range from 0.1% to 2% (**Illumina, 2017; van der Valk et al., 2023**). Collectively, these observations strongly support that the recovered viral genomes represent bona fide mycoviruses associated with *T. thlaspeos*, rather than contaminants originating from plant material. Consistent with this interpretation, viral genome coverage in fungal libraries ranged from 4× to 404×, whereas coverage values in plant libraries remained extremely low (0–0.7×). These coverage values are well above those typically considered sufficient for reliable viral genome assembly from RNA-seq data (e.g., **Gilbert et al., 2019**).

It is well established that the relative abundance of aminoacyl-tRNAs in the host cytoplasm imposes translational selection on synonymous codon usage in viral genomes (**Bulmer, 1987; Kumar et al., 2018; Chen et al., 2020**). As a consequence, viral coding sequences often converge toward host codon usage preferences, reflecting adaptation to the host translational machinery and optimization of translation efficiency. Consistent with this expectation, cosine distances between viral and host RSCU profiles were very small (0.0084–0.0109), indicating highly similar codon preferences. Although such similarity may partly reflect shared mutational biases, it is also compatible with evolutionary adaptation of viral coding sequences to the codon usage bias of *T. thlaspeos*. Similar patterns have been reported in other host-associated viral communities and are often interpreted as optimization for abundant tRNA species and efficient translation (**Bahir et al., 2009**).

At the same time, our RSCU analysis revealed subtle but detectable differences among viral strains. Such variation may reflect the combined influence of neutral evolutionary processes and mutational pressures contributing to nucleotide divergence (**Jacquat et al., 2024**). RNA viruses are known to exhibit high mutation rates, typically on the order of 10⁻⁴–10⁻³ substitutions per nucleotide site per replication cycle, largely due to the error-prone replication of RdRps (**Drake and Holland, 1999**). Experimental analyses of mutation accumulation in Fusarium graminearum virus 1–4 (FgV1–4) have demonstrated that viral evolutionary trajectories could differ with respect to hosts and also with respect to co-infecting viruses (**Heo et al., 2020**). We identified moderate nucleotide-level variability among virus strains shared by mating types a1b1 and a2b2. However, viral proteins are highly conserved at the amino-acid level, which suggests that purifying selection eliminates most nonsynonymous mutations. Such a pattern, substantial synonymous divergence coupled with strong conservation of protein sequences, is commonly observed in RNA viruses, reflecting the need to maintain essential structural and enzymatic functions (**Duffy et al., 2008**).

The eight viral elements, ranging from 4,480 to 5,296 nt, are monosegmented and bicistronic, with two overlapping ORFs encoding a CP and an RdRp. These viruses are predicted to employ a translational recoding strategy whereby the RdRp is produced as the C-terminal domain of a CP–RdRp fusion protein via −1 PRF. Notably, differences were observed between eimeriaviruses and totiviruses. In all five totiviruses, a conserved slippery heptanucleotide motif of the form NNN_WWW_H was identified, with two closely related variants (GGA_TTT_T and GGG_TTT_T) differing by a single nucleotide. This pattern is consistent with previously described totivirus slippery sequences, where GG(A/G)TTT_T represents a frequent consensus motif (**figure 5B**). It is well established that such heptanucleotide motifs, together with downstream RNA secondary structures such as pseudoknots or stem-loops, promote efficient −1 ribosomal frameshifting (**Dinman et al., 1991; Dinman, 2012**).

Minus-one translational recoding typically requires a stimulatory RNA structure located within ∼12 nucleotides downstream of the slippery site, including pseudoknot or other stimulatory structures as hairpin variants (**Dinman, 2012; Chang and Wen, 2021**). In the present study, multiple computational approaches (KnotInFrame, ProbKnot, and IPknot) identified candidate pseudoknotted structures downstream of the predicted slippery motifs in all five totivirus genomes. Comparative thermodynamic analyses, based on normalized folding free energies and the relative Δrel metric, indicated that these pseudoknotted conformations were generally less stable than the corresponding pseudoknot-free (nested) structures, suggesting that they do not represent the global minimum free-energy conformation.

However, increasing evidence indicates that −1 PRF efficiency is not solely determined by thermodynamic stability, but also by the structural dynamics of the RNA (**Chang and Wen, 2021**). In particular, the propensity of stimulatory elements to adopt alternative conformations, referred to as conformational plasticity, has been shown to positively correlate with frameshifting efficiency (**Chang and Wen, 2021**). Multiple experimental studies show that RNAs sampling alternative or metastable folds promote frameshifting, whereas pseudoknot mechanical resistance to unfolding does not primarily determine frameshifting efficiency (**Ritchie et al., 2012; Halma et al., 2012**). In this context, the lower thermodynamic stability of the predicted pseudoknots in *T. thlaspeos* totiviruses may reflect a dynamic folding landscape in which transient or alternative conformations are functionally relevant. Such conformational heterogeneity could modulate −1 frameshifting efficiency by controlling the temporal availability and mechanical resistance of the stimulatory structure during ribosome translocation. Consequently, these metastable pseudoknots may contribute to maintaining a regulated, low-frequency production of the CP–RdRp fusion protein relative to CP alone, consistent with models of stoichiometric balance required for proper totivirus particle assembly.

The three eimeriavirus genome sequences lack a recognizable GG(A/G)TTTT-like slippery motif, suggesting an evolutionary divergence in translational strategy relative to totiviruses. Instead, they comprise two partially overlapping ORFs separated by a highly conserved tetranucleotide ATGA junction. This genomic organization is consistent with a termination–reinitiation mechanism similar to that described in Helminthosporium victoriae virus 190S, a closely related virus in which this strategy has been experimentally validated (**Li et al. 2011**). However, the molecular determinants underlying this process in eimeriaviruses remain unresolved and warrant direct experimental investigation.

### Biological insights

The differential virome composition between strains LF1 (mating type a1b1) and LF2 (mating type a2b2) warrants significant attention. The complete absence of TtEV2, TtEV3, and TtTV2 from LF1, representing the loss of two of the three *Eimeriavirus* species and one of the five *Totivirus* species, suggest a pattern of strain-specific virome architecture. Several non-mutually exclusive hypotheses could account for this observation. First, independent virus acquisition or loss during the evolutionary or laboratory history of these strains could generate the observed pattern. Second, virus–virus interactions, such as superinfection exclusion or mutual dependence among co-infecting viruses, may restrict the range of viable virome configurations. Third, incompatibility between certain viruses and the LF1 genetic background could limit viral establishment or persistence. The recent demonstration by **Kuroki et al. (2023)** that mycoviral effects on fungal hosts are strongly modulated by host strain genotype lends support to this hypothesis. In natural populations of *T. thlaspeos*, horizontal transmission through hyphal anastomosis would be expected to promote partial homogenization of virome composition among compatible strains. The apparent strain-associated infection pattern observed here, however, suggests that viral establishment may be constrained by host genetic background, vegetative compatibility barriers, or lineage-specific virus–host adaptation, ultimately leading to segregation of viral genotypes among host lineages.

The observed hierarchy of viral transcript abundance, characterized by one or two dominant viruses coexisting with several lower-abundance components, is consistent with patterns reported in other complex mycoviromes (**Osaki et al., 2016; Gilbert et al., 2019**). As an extreme example, *Diplodia seriata* (*Botryosphaeriaceae*) was found to harbor eight mycoviruses belong to seven different viral families, including totivirus-like viruses (**Khan et al., 2022**). Such structured viral communities may reflect stable coexistence dynamics within the fungal host rather than random viral accumulation. Co-infecting mycoviruses can engage in a wide range of interactions. Some are synergistic, enhancing replication and promoting vertical and horizontal transmission, even across vegetative incompatibility barriers (**Sun et al., 2006; Kashif et al., 2019**), while others are antagonistic and limit co-infection (**Heo et al., 2020**). Notably, many of these experimental observations derive from controlled or artificial infection systems. For example, antagonistic interactions between FgV1 and FgV2 were demonstrated in *Fusarium graminearum* strains naturally infected with FgV1 and subsequently experimentally co-infected with FgV2 originating from *F. asiaticum* (**Heo et al., 2020**). While such studies provide valuable mechanistic insights, the extent to which these interactions reflect natural co-infection dynamics remains uncertain. In contrast, studies in more natural host–virus systems have yielded mixed outcomes. For instance, **Li et al. (2022)** showed that infection by AaCV1-QY2 alone significantly altered host morphology and reduced virulence in *Alternaria alternata*. However, comparable levels of hypovirulence were observed in the same *A. alternata* strain with double infection, suggesting that a single dominant virus may drive the phenotype, while additional co-infecting viruses may have limited or context-dependent effects (**Li et al., 2022**).

Moreover, differences in the composition or relative abundance of these viruses may have functional consequences for the host. In this context, recent experimental work by **Kuroki et al. (2023)** demonstrated that mycoviral effects on host phenotype and transcriptome can be highly strain-dependent in *Aspergillus flavus*, where even minimal sequence variation in the viral genome, sometimes as few as three nucleotide substitutions, was sufficient to elicit distinct host responses. In contrast, studies in *U. virens* suggest a more generalist interaction, in which mycoviruses are able to infect multiple host isolates independently of their genetic background (**Zhong et al., 2017**). Together, these observations highlight that virus–host interactions in fungi can range from highly genotype-specific to broadly permissive, while interactions among co-infecting mycoviruses are similarly context-dependent, spanning synergistic, neutral, or antagonistic outcomes. Collectively, this underscores the need for system-specific investigation of mycoviral ecology and evolution within natural host–virus frameworks.

The elevated TtEV1 RPKM in LF1 relative to LF2 (28.29 vs. 13.27 ± 7.34) is an intriguing observation that invites further investigation. One plausible interpretation is that TtEV1 accumulation increases in the absence of competing viral species, a dynamic consistent with the ecological concept of competitive release. In multi-virus fungal systems, resource competition for host replication machinery has been suggested as a factor limiting individual viral loads (**Thapa and Roossinck, 2019**).

From a biological perspective, the fact that both strains appear to tolerate their respective viral complements under culture conditions could suggest that these viruses are either neutral or beneficial to fungal fitness, at least under the growth conditions tested. However, phenotypic effects that manifest only during plant infection cannot be excluded.

## Conclusion and future directions

In conclusion, our work unveils the previously unexplored viral landscape of the biotrophic pathogen *T. thlaspeos*, expanding the known diversity of the order *Ghabrivirales*. By identifying five novel totiviruses and three novel eimeriaviruses, we demonstrate the power of retrospective data mining for virus discovery in agriculturally significant fungi. This study serves as a critical starting point for deciphering the biological relevance of the *T. thlaspeos* virome within the context of plant pathology and the development of virus-mediated biocontrol strategies.

Several methodological caveats should be acknowledged. The LF1 dataset derives from a single replicate with single-end, shorter reads, which limits both statistical power and assembly completeness. The plant control analysis, while strongly supporting fungal specificity, relies on read-count thresholds. Incorporation of formal statistical models (e.g., binomial or Poisson frameworks accounting for expected index-hopping rates) would further strengthen these conclusions. Moreover, as the present data derive from axenic mycelial cultures, the virome composition during active plant infection may differ from that observed under in vitro conditions.

This study establishes *T. thlaspeos* as a host of a complex RNA virome comprising eight novel viruses from two distinct families. The observed strain-specific differences in virome composition represent, to our knowledge, one of the first documented cases of intraspecific virome variation in smut fungi, supporting the idea that host genetic background plays an important role in shaping mycoviral communities. Key priorities for future research include the additional biological replicates and field isolates to assess virome prevalence and stability; and strand-specific RNA-seq to evaluate active viral replication; characterization of phenotypic effects through virus-cured isogenic lines Experimental validation should also include RT-PCR amplification of viral RNA from independent biological samples to confirm the presence of these viruses in *T. thlaspeos*. In addition, rapid amplification of cDNA ends (RACE) approaches will be necessary to resolve the complete 5′ and 3′ genome termini, thereby improving genome annotation and validating the inferred genome architectures. Finally, it is important to recognize that RNA secondary structure predictions used to explore potential regulatory elements associated with translational recoding in both totivirus- and eimeriavirus-like genomes have inherent limitations. RNA molecules do not adopt a single static conformation but instead exist as dynamic ensembles of alternative structures in thermodynamic equilibrium. Consequently, the predicted conformations represent the most probable states under the applied model, yet may not fully reflect the structural landscape in vivo (**Schuster et al., 1994; Vandivier et al., 2016; Tahi et al., 2017**). Experimental validation will therefore be required to determine the functional impact of these predicted structures on translational recoding efficiency, for example through reporter-based assays combined with RNA structure probing approaches, particularly to distinguish between −1 PRF-dependent mechanisms in totiviruses and potential termination–reinitiation strategies in eimeriaviruses.

## Data Availability

The genomes of the identified *T. thlaspeos* viruses have been submitted to NCBI GenBank and are available as **Supplementary Material 1** of this submission.

## Acknowledgments

We thank the authors of Courville et al. (2019) for generating the RNA-seq datasets that enabled this comparative analysis.

## Bibliography

Arias, R. S., Dobbs, J. T., Orner, V. A., Conforto, E. C., Rago, A. M., Cazon, L. I., … and Massa, A. N. (2025). First metagenome-and metatranscriptome dataset of Thecaphora frezzii teliospores, assembly and annotation of a new bacterial genome. Data in Brief, 61, 111779. DOI: 10.1016/j.dib.2025.111779.

Arias, S. L., Mary, V. S., Velez, P. A., Rodriguez, M. G., Otaiza-González, S. N., and Theumer, M. G. (2021). Where does the peanut smut pathogen, Thecaphora frezii, fit in the spectrum of smut diseases?. Plant disease, 105(9), 2268–2280. DOI: 10.1094/PDIS-11-20-2438-FE

Bahir, I., Fromer, M., Prat, Y., and Linial, M. (2009). Viral adaptation to host: a proteome-based analysis of codon usage and amino acid preferences. Molecular systems biology, 5(1), MSB200971. DOI: 10.1038/msb.2009.71.

Bejerman, N., Dietzgen, R. G., and Debat, H. (2025). Expanding the known nucleorhabdovirus world: the final chapter in a trilogy exploring the hidden diversity of plant-associated rhabdoviruses. Virology, 110699. DOI: 10.1016/j.virol.2025.110699

Bellaousov, S., and Mathews, D. H. (2010). ProbKnot: fast prediction of RNA secondary structure including pseudoknots. Rna, 16(10), 1870–1880. DOI: 10.1261/rna.2125310.

Bulmer, M. (1987). Coevolution of codon usage and transfer RNA abundance. Nature, 325(6106), 728–730. DOI: 10.1038/325728a0.

Camacho, C., Coulouris, G., Avagyan, V., Ma, N., Papadopoulos, J., Bealer, K., and Madden, T. L. (2009). BLAST+: architecture and applications. BMC bioinformatics, 10(1), 421. DOI: 10.1186/1471-2105-10-421

Chan, C. Y., and Ding, Y. (2008). Boltzmann ensemble features of RNA secondary structures: a comparative analysis of biological RNA sequences and random shuffles. Journal of mathematical biology, 56(1-2), 93–105. DOI: 10.1007/s00285-007-0129-z

Chang, K. C., and Wen, J. D. (2021). Programmed− 1 ribosomal frameshifting from the perspective of the conformational dynamics of mRNA and ribosomes. Computational and Structural Biotechnology Journal, 19, 3580–3588. DOI: 10.1016/j.csbj.2021.06.015.

Chen, F., Wu, P., Deng, S., Zhang, H., Hou, Y., Hu, Z., … and Yang, J. R. (2020). Dissimilation of synonymous codon usage bias in virus–host coevolution due to translational selection. Nature ecology and evolution, 4(4), 589–600. DOI: 10.1038/s41559-020-1124-7

Chen, S., Cao, L., Huang, Q., Qian, Y., and Zhou, X. (2016). The complete genome sequence of a novel maize-associated totivirus. Archives of virology, 161(2), 487–490. DOI: 10.1007/s00705-015-2657-y

Courville, K. J., Frantzeskakis, L., Gul, S., Haeger, N., Kellner, R., Heßler, N., … and Göhre, V. (2019). Smut infection of perennial hosts: the genome and the transcriptome of the Brassicaceae smut fungus Thecaphora thlaspeos reveal functionally conserved and novel effectors. New Phytologist, 222(3), 1474–1492. DOI: 10.1111/nph.15692

Debat, H., Farrher, E. S., and Bejerman, N. (2024). Insights into the RNA virome of the corn leafhopper Dalbulus maidis, a major emergent threat of Maize in Latin America. Viruses, 16(10), 1583. DOI: 10.3390/v17010095

Debat, H., Gomez-Talquenca, S., and Bejerman, N. (2025b). RNA virus discovery sheds light on the virome of a major vineyard pest, the European grapevine moth (Lobesia botrana). Viruses, 17(1), 95.

Debat, H., Paolinelli, M., Escoriaza, G., Garcia-Lampasona, S., Gomez-Talquenca, S., and Bejerman, N. (2025a). Grapevine holobiome metatranscriptomics provides a glimpse into the wood mycovirome. Virology, 610, 110604. DOI: 10.1016/j.virol.2025.110604

Díaz, M. S., Soria, N. W., Figueroa, A. C., Yang, P., Badariotti, E. H., Alasino, V. R., … and Beltramo, D. M. (2024). Transcriptional study of genes involved in the passage from teliospore to hyphae stage in the fungus Thecaphora frezii, the causal agent of peanut smut. Revista argentina de microbiología, 56(2), 175–186. DOI: 10.1016/j.ram.2023.10.002

Dinman, J. D. (2012). Mechanisms and implications of programmed translational frameshifting. Wiley Interdisciplinary Reviews: RNA, 3(5), 661–673. DOI: 10.1002/wrna.1126

Dinman, J. D., Icho, T., and Wickner, R. B. (1991). A-1 ribosomal frameshift in a double-stranded RNA virus of yeast forms a gag-pol fusion protein. Proceedings of the National Academy of Sciences, 88(1), 174–178. DOI: 10.1073/pnas.88.1.174

Duffy, S., Shackelton, L. A., and Holmes, E. C. (2008). Rates of evolutionary change in viruses: patterns and determinants. Nature Reviews Genetics, 9(4), 267–276. DOI: 10.1038/nrg2323.

Feschotte, C., and Gilbert, C. (2012). Endogenous viruses: insights into viral evolution and impact on host biology. Nature Reviews Genetics, 13(4), 283–296. DOI: 10.1038/nrg3199

Frantzeskakis, L., Courville, K. J., Plücker, L., Kellner, R., Kruse, J., Brachmann, A., … and Göhre, V. (2017). The plant-dependent life cycle of Thecaphora thlaspeos: a smut fungus adapted to Brassicaceae. Molecular Plant-Microbe Interactions, 30(4), 271–282. DOI: 10.1094/MPMI-08-16-0164-R

García-Pedrajas, M. D., Cañizares, M. C., Sarmiento-Villamil, J. L., Jacquat, A. G., and Dambolena, J. S. (2019). Mycoviruses in biological control: From basic research to field implementation. Phytopathology, 109(11), 1828–1839. DOI: 10.1094/PHYTO-05-19-0166-RVW

Ghabrial, S. A., Castón, J. R., Jiang, D., Nibert, M. L., and Suzuki, N. (2015). 50-plus years of fungal viruses. Virology, 479-480, 356–368. DOI: 10.1016/j.virol.2015.02.034

Gilbert, K. B., Holcomb, E. E., Allscheid, R. L., and Carrington, J. C. (2019). Hiding in plain sight: New virus genomes discovered via a systematic analysis of fungal public transcriptomes. PloS one, 14(7), e0219207. DOI: 10.1371/journal.pone.0219207

Gruber, A. R., Lorenz, R., Bernhart, S. H., Neuböck, R., and Hofacker, I. L. (2008). The vienna RNA websuite. Nucleic acids research, 36(suppl_2), W70-W74. DOI: 10.1093/nar/gkn188.

Halma, M. T., Ritchie, D. B., Cappellano, T. R., Neupane, K., and Woodside, M. T. (2019). Complex dynamics under tension in a high-efficiency frameshift stimulatory structure. Proceedings of the National Academy of Sciences, 116(39), 19500–19505. DOI: 10.1073/pnas.1905258116.

Heo, J. I., Yu, J., Choi, H., and Kim, K. H. (2020). The signatures of natural selection and molecular evolution in Fusarium graminearum Virus 1. Frontiers in Microbiology, 11, 600775. DOI: 10.3389/fmicb.2020.600775.

Hoang, D. T., Chernomor, O., Von Haeseler, A., Minh, B. Q., and Vinh, L. S. (2018). UFBoot2: improving the ultrafast bootstrap approximation. Molecular biology and evolution, 35(2), 518–522. DOI:10.1093/molbev/msx281

Hough, B., Steenkamp, E., Wingfield, B., and Read, D. (2023). Fungal viruses unveiled: A comprehensive review of mycoviruses. Viruses, 15(5), 1202. 10.3390/v15051202

ICTV (International Committee on Taxonomy of Viruses). 2024. Proposal 2023.015F.Ghabrivirales_reorg. Ratified in Master Species List (MSL) #39, Release v2 (October 30, 2024). Available at: https://ictv.global/.

Illumina, Inc. (2017). Effects of index misassignment on multiplexing and downstream analysis (White paper No. 770-2017-004). Illumina https://assets.illumina.com/content/dam/illumina-marketing/documents/products/whitepapers/index-hopping-white-paper-770-2017-004.pdf (acceded at March 8, 2026)

Jacquat, A. G., Theumer, M. G., and Dambolena, J. S. (2024). Selective and non-selective evolutionary signatures found in the simplest replicative biological entities. Journal of Evolutionary Biology, 37(8), 862–876. DOI: 10.1093/jeb/voae070

Jiang, Y., Zhang, T., Luo, C., Jiang, D., Li, G., Li, Q., … and Huang, J. (2015). Prevalence and diversity of mycoviruses infecting the plant pathogen Ustilaginoidea virens. Virus Research, 195, 47–56. DOI: 10.1016/j.virusres.2014.08.022

Jo, Y., Choi, H., Chu, H., and Cho, W. K. (2022). Unveiling mycoviromes using fungal transcriptomes. International Journal of Molecular Sciences, 23(18), 10926. DOI: 10.3390/ijms231810926.

Johnson, P. Z., and Simon, A. E. (2023). RNAcanvas: interactive drawing and exploration of nucleic acid structures. Nucleic acids research, 51(W1), W501–W508. DOI: 10.1093/nar/gkad302

Kalyaanamoorthy, S., Minh, B. Q., Wong, T. K., Von Haeseler, A., and Jermiin, L. S. (2017). ModelFinder: fast model selection for accurate phylogenetic estimates. Nature methods, 14(6), 587–589. DOI: 10.1038/nmeth.4285.

Kashif, M., Jurvansuu, J., Vainio, E. J., and Hantula, J. (2019). Alphapartitiviruses of Heterobasidion wood decay fungi affect each other’s transmission and host growth. Frontiers in Cellular and Infection Microbiology, 9, 64.

Katoh, K., and Standley, D. M. (2013). MAFFT multiple sequence alignment software version 7: Improvements in performance and usability. Molecular Biology and Evolution, 30(4), 772–780. DOI: 10.1093/molbev/mst010

Katoh, K., Rozewicki, J., and Yamada, K. D. (2019). MAFFT online service: Multiple sequence alignment, interactive sequence choice and visualization. Briefings in Bioinformatics, 20(4), 1160–1166. DOI: 10.1093/bib/bbx108

Khan, H. A., Telengech, P., Kondo, H., Bhatti, M. F., and Suzuki, N. (2022). Mycovirus hunting revealed the presence of diverse viruses in a single isolate of the phytopathogenic fungus Diplodia seriata from Pakistan. Frontiers in Cellular and Infection Microbiology, 12, 913619. DOI: 10.3389/fcimb.2022.913619

Kondo, H., Kanematsu, S., and Suzuki, N. (2013). Viruses of the white root rot fungus, Rosellinia necatrix. Advances in virus research, 86, 177–214. DOI: 10.1016/B978-0-12-394315-6.00007-6

Kronstad, J. W. (1996). Pathogenesis and sexual development of the smut fungi. Plant-microbe interactions, 1, 141–186. DOI: 10.1007/978-1-4613-1213-0_5

Kumar, N., Kulkarni, D. D., Lee, B., Kaushik, R., Bhatia, S., Sood, R., … and Singh, V. P. (2018a). Evolution of codon usage bias in Henipaviruses is governed by natural selection and is host-specific. Viruses, 10(11), 604. DOI: 10.3390/v10110604.

Kuroki, M., Yaguchi, T., Urayama, S. I., and Hagiwara, D. (2023). Experimental verification of strain-dependent relationship between mycovirus and its fungal host. Iscience, 26(8). DOI: 10.1016/j.isci.2023.107337.

Langmead, B., and Salzberg, S. L. (2012). Fast gapped-read alignment with Bowtie 2. Nature methods, 9(4), 357–359. DOI: 10.1038/nmeth.1923

Li, B., Cao, Y., Ji, Z., Zhang, J., Meng, X., Dai, P., … and Wang, Y. (2022). Coinfection of two mycoviruses confers hypovirulence and reduces the production of mycotoxin alternariol in Alternaria alternata f. sp. mali. Frontiers in Microbiology, 13, 910712. DOI: 10.3389/fmicb.2022.910712.

Li, W., and Godzik, A. (2006). Cd-hit: a fast program for clustering and comparing large sets of protein or nucleotide sequences. Bioinformatics, 22(13), 1658–1659. DOI: DOI: 10.1093/bioinformatics/btl158

Liu, M., Ni, Y., Wang, J., Liu, X., Zhao, H., Zhao, X., … and Miao, H. (2025). Diverse, novel mycoviruses coinfecting the phytopathogenic fungus Corynespora cassiicola from Sesamum indicum. Frontiers in Cellular and Infection Microbiology, 15, 1704628. DOI: 10.3389/fcimb.2025.1704628

Lorenz, R., Bernhart, S. H., Höner zu Siederdissen, C., Tafer, H., Flamm, C., Stadler, P. F., and Hofacker, I. L. (2011). ViennaRNA Package 2.0. Algorithms for molecular biology, 6, 1–14. DOI: 10.1186/1748-7188-6-26.

Marchler-Bauer, A., and Bryant, S. H. (2004). CD-Search: protein domain annotations on the fly. Nucleic acids research, 32(suppl_2), W327-W331. Doi: 10.1093/nar/gkh454

Minh, B. Q., Schmidt, H. A., Chernomor, O., Schrempf, D., Woodhams, M. D., Von Haeseler, A., and Lanfear, R. (2020). IQ-TREE 2: new models and efficient methods for phylogenetic inference in the genomic era. Molecular biology and evolution, 37(5), 1530–1534. DOI: 10.1093/molbev/msaa015.

Nibert, M. L., Debat, H. J., Manny, A. R., Grigoriev, I. V., and De Fine Licht, H. H. (2019). Mitovirus and mitochondrial coding sequences from basal fungus Entomophthora muscae. Viruses, 11(4), 351. DOI: 10.3390/v11040351

Osaki, H., Sasaki, A., Nomiyama, K., and Tomioka, K. (2016). Multiple virus infection in a single strain of Fusarium poae shown by deep sequencing. Virus Genes, 52(6), 835–847. DOI: 10.1007/s11262-016-1379-x.

Plücker, L., Bösch, K., Geißl, L., Hoffmann, P., and Göhre, V. (2021). Genetic manipulation of the Brassicaceae smut fungus Thecaphora thlaspeos. Journal of Fungi, 7(1), 38. DOI: 10.3390/jof7010038

Reuter, J. S., and Mathews, D. H. (2010). RNAstructure: software for RNA secondary structure prediction and analysis. BMC bioinformatics, 11(1), 129. DOI: 10.1186/1471-2105-11-129

Ritchie, D. B., Foster, D. A., and Woodside, M. T. (2012). Programmed− 1 frameshifting efficiency correlates with RNA pseudoknot conformational plasticity, not resistance to mechanical unfolding. Proceedings of the National Academy of Sciences, 109(40), 16167–16172. DOI: 10.1073/pnas.1204114109.

Sato, K., and Kato, Y. (2022). Prediction of RNA secondary structure including pseudoknots for long sequences. Briefings in Bioinformatics, 23(1), bbab395. DOI: 10.1093/bib/bbab395

Sato, K., Kato, Y., Hamada, M., Akutsu, T., and Asai, K. (2011). IPknot: fast and accurate prediction of RNA secondary structures with pseudoknots using integer programming. Bioinformatics, 27(13), i85–i93. DOI: 10.1093/bioinformatics/btr215

Schuster, P., Fontana, W., Stadler, P. F., and Hofacker, I. L. (1994). From sequences to shapes and back: a case study in RNA secondary structures. Proceedings of the Royal Society of London. Series B: Biological Sciences, 255(1344), 279–284. DOI: 10.1098/rspb.1994.0040.

Sun, L., Nuss, D. L., and Suzuki, N. (2006). Synergism between a mycoreovirus and a hypovirus mediated by the papain-like protease p29 of the prototypic hypovirus CHV1-EP713. Journal of general virology, 87(12), 3703–3714. DOI: doi: 10.1099/vir.0.82213-0.

Sutela, S., Forgia, M., Vainio, E. J., Chiapello, M., Daghino, S., Vallino, M., … and Turina, M. (2020). The virome from a collection of endomycorrhizal fungi reveals new viral taxa with unprecedented genome organization. Virus Evolution, 6(2), veaa076. DOI: 10.1093/ve/veaa076

Tahi, F., Du T. Tran, V., and Boucheham, A. (2017). In silico prediction of RNA secondary structure. Promoter Associated RNA: Methods and Protocols, 145-168. DOI: 10.1007/978-1-4939-6716-2_7.

Thapa, V., and Roossinck, M. J. (2019). Determinants of coinfection in the mycoviruses. Frontiers in cellular and infection microbiology, 9, 169. DOI: /10.3389/fcimb.2019.00169.

Theis, C., Reeder, J., and Giegerich, R. (2008). KnotInFrame: prediction of− 1 ribosomal frameshift events. Nucleic acids research, 36(18), 6013–6020. DOI: 10.1093/nar/gkn578

Trifinopoulos, J., Nguyen, L. T., von Haeseler, A., and Minh, B. Q. (2016). W-IQ-TREE: a fast online phylogenetic tool for maximum likelihood analysis. Nucleic acids research, 44(W1), W232–W235.

van der Valk, T., Vezzi, F., Ormestad, M., Dalén, L., and Guschanski, K. (2020). Index hopping on the Illumina HiseqX platform and its consequences for ancient DNA studies. Molecular ecology resources, 20(5), 1171–1181. DOI: 10.1111/1755-0998.13009.

Vandivier, L. E., Anderson, S. J., Foley, S. W., and Gregory, B. D. (2016). The conservation and function of RNA secondary structure in plants. Annual review of plant biology, 67, 463–488. DOI:10.1146/annurev-arplant-043015-111754.

Wickner, R. B., Fujimura, T., and Esteban, R. (2013). Viruses and prions of Saccharomyces cerevisiae. Advances in virus research, 86, 1–36. DOI: 10.1016/B978-0-12-394315-6.00001-5

Zhong, J., Cheng, C. Y., Gao, B. D., Zhou, Q., and Zhu, H. J. (2017). Mycoviruses in the plant pathogen Ustilaginoidea virens are not correlated with the genetic backgrounds of its hosts. International Journal of Molecular Sciences, 18(5), 963. DOI: 10.3390/ijms18050963.

Zuber, J., and Mathews, D. H. (2019). Estimating uncertainty in predicted folding free energy changes of RNA secondary structures. RNA, 25(6), 747–754. doi: 10.1261/rna.069203.118

