## Supplementary Material 1 for "Novel RNA viruses reveal a complex mycovirome in the smut fungus *Thecaphora thlaspeos*"

>Thecaphora_thlaspeos_assocoated_totivirus_1

CGAAATTATCCCCGTGATGCTTTCGAATCTTTACTCTACGCTCGGGTCGTTCGCGGCCATAAAACACGCTGCCCTTTCCGCCTTTGGGGGCTTTACCACTAAAAGGCGCATCAAAATCGAAGTTCACGATGGGGCCAATAAGGCCAAACGTGATTTTGTGGCTGCAGAAAGCTACACCGCCGTCGGCGTCGCCGACGTTGAATTCGGCCCCCCTCTGAACGATGTCCGCGGATACGATACCGCCATCATCGATGAGGCCGGGGTGCTGACACCCGAACTATTTCTGGCGCGCATGCGCACGATGGGCGGCGTTTCCCGCCCCACGCTCACGGACATGGCTGTCGCACCGGAGCAGTACATCAGGGATGACTCCCAGATTGCTCTGCTGGTGTGTCTGCTTTCGACGTACTATGTTCTTTCGGCTGACGATCAGGCGCGTAAGTGCTTGATTTCGGATTACAATGATGGGCATGTGCGTATCAAAGGATCTTGTCTGGCGCCGGACGTTAAACTGGCCGACAAATGGTGGTATAACGACCTTATCGATATGGAAAGCTGGCAGCAGGCCAATTTTCAGATAGCGCCGGGTTACCCCGTGCTACAGATGGGACGTATGTCTGCAGCCGACGCTGCTATTGTCGTCACTCACTTGGCTGGCCGTAAGAAAACGACCAACCTGTTGTTTGACTTGGAAATCCCGGCCTTAGCCGATCACGTCGCCATCGCGGGGCTCACCCCCCCCGATGAGGTTCCCGAGAACGTCACCGCTGCCAAAGTCTTGAGGGCTCTCAACAAGTACGTTGAAGTAAACAGGCTTCATGCAACCTTCGAGGCTGCACTGGGCATATATGCTCAGCTGGCTCTCAACCCCGTCCCCGATACGGCCGAGGCTCTCGTCTGGCTGAATCGGAAGAAGACAGTTGTGCTTCCGTCGTTTGACGCCTACCGTGGCGCCTACCGCGTACTGACGCTGGAGAAGCCGTACGGCTTCTCGTCAGCAGTAACTGAGACGTGGAAAGCGTGGCACGAGCGGGGTCTGGCGATTGCCGTCCCGTTCGGCGTTTTCAACACTTTGTCGTTATGGGGCCGGTACTTGGTTGAAGGACGGTTTTACCACAGCGACGCCGAAATCGATCGTCTGGGCACTCTCGAGTTTGCTGAGGCACCGGCAATGGCCTACTACGCGTACGCTGCCTTGATCACCGGGACGGACGTGTATTGTGCTGTCCCGCCGGGTTTCGGGCTGCACTACGCTGCATGGGACTTCGCTGAAGGAGTACGTCTACCGGTGGACGTGCTCGATAACGCGGTGGCCGGATACAACATCGAACTCGACACTGCGGGCAGCATGCTCGTGGTCAAACAGTTGCCGCCGCCGTGCTCCCCCGTCAGTGTGATGGGACGATTGCCCCGAGAGGGAAGTTTTAACGCTCTCTCGGATGAGATGACTGTGACGATGCGGCACATCAGCGACGGCTGGGTGCTAGACAACGTCAACGACGCTTGGAAGATGGCGATTGCAACCCGGCTCTTGGGTTACAACGCCGTCATGGAGTACAACGGGCAAGAGAAGATTGCGAACTGGGCCTCGAACCAGACGTCAATCCCTTCGCGCCCGTTTTTCGTGGGTGGGCCTGACCGTCAGACGGTCAAGCTCAAGTCGATATATGAGAGGAAAAAAGTCTTCTCGCCGCTGCCATCGCTTGCCGGGGATTCGTCTACCTGGAAGATCAATATCATGGTATCAGACGACTTCAGCATCGGAGGGCCTAACGCCCATCGCTTCGGTGTTGTCGGCACATTCGTTAGGCCGAGGGTCGTCGACCCTCAATTGTACACGCCTCAGGCCCACACCCTGACGCAGATTCCCGTGGTTATCAGGGCGGTCCGCCGTGCTGAGGCCACGATGGCGGGTTTTCAAGTGGCGGACCTGACTACCGCCCACCATCCCCCACCCATACAACGCGACCCGGTTGCAGAGCAGCCAGAGGAAGCGGGGGAGACCGCACCGGATGGAGGGGAATAAGATTCTCCAAATCCGAATTCATGCCCCTGCTTCTGCGAGTGACACAGAACGGGTACGCCCTAGCGGACGTAGAGGATGCTACCTTGGCATTAGTACATGTAGAGGCCAGTTCTAATAGAAATAGCGGAACGTGGCGGCACAACGTCAAGGGTGAGTCGGTTTATCTGACGCATGTGCCATGGCGTGGGCTCCAGTTGTGTTATATGGACACTTCTCAGGCCACCGCGGCATCAGTGGCAGTAAGCAGAGTGGCGGGCCTCTCCTTACTGGCTGATGAGATTTTCGTCGACTACGGTGCTAACGTGCCGTTCCTGGATCTGACCAACGGTAACACATATCGGCCGCCGGGCAACACCAACCGGGCGGTGCGAGCTATCGTTGTATGTCGGTCAAGCGACCGGCCTGATGAAGCGCTGGCTTCAGCAGGTTATGGACCTTGGCTTGACACAAAGGTGACTGGGCGACATCACACTTGGTTTACGCCTGGTGAGATCGTTGCTAGCGCCCCAAACGCTTTTTCGCGACAGCTGGCTGTAGCGCTAGAGCGTATGCGGGCTTTCCCAAGTAGTAGGGCGTTTGTAGCGGGTTATCTACTGTGGGCGATCTCTGCAGATAGCAGAGTCGTAGACCACATTAACCACTGCGACCGCTTGTGGAGCGTCAACACTCTGGAAGGCTGGGCGTCGCGCGCAAAGGCGTTGACGTCCGCTTATAAGTTGCACCACCACTACGTTGATCTGCCGCTCGAGCAGGCGTTCGAGCTGCAGGTGTTGGTGAATAGAGGAGTGGGAGAAGTGGACTGGTTCGCGGAGAAGGACCACCGTGTTAACCCTGTGTTGGCAGAGTTCAAATACAAGGACGTGTTTGATCAGGCGGTGCAGATCTTCAGGGGAGGTGCAGCCCAGGGATACAAGTATAAAGAGGAGACTTGGGGCAACTTCTGGCGGCGTAGGTGGGCCTCAGTGCCTGGTGGTAGCGTACATAGTCAGCATGCGGACGACACTCCTTACATTGAGAAAGAACAGTCCAAGAGGACGAAACACTACACACTCATTAAGATGTCTGACTCTAAGCTACGCAGGTGGACGGGCCGCGCGCCGGAGATTGCAGCATGGAAGTCTGTGAAGTACGAATGGGCCAAGAATAGGGCTATATACGGCACAGACCTTACTTCGTACTTGCTTACGGACTTTGCCATGCCCAACGTTGAGGAGTCGCTGACTCACATGTTCCCGGTGGGTAAGAAGGCCGACGAGGCGTACGTGACGAGACGAGTCGAGCTCGCGGGCGAGGGCGGGATCGCATTCTGCTTCGATTATGAGGACTTCAATAGTCAACATTCGATCACGTCACAGCTCGCCGTGGTCGCAGCGTACCAGGACGTCTTCAGCAACCAGATGTCTGCGTCGCAGCAGGAAGCAATGGAGTGGGTTGTGCAGAGTGTGAAACGGCAGACCGTGGCTGGGGACGATGGCTACGGAACGAACGGGACTTTGCTGAGCGGATGGCGCTTGACCACACTCGTTAATACGGTACTTAACTATGTGTACCTTAAAATTTCTGGCGCCACCGACTGCTTCCGTGATTCGCTTCACAATGGCGACGATGTACTCGCTTACACTGATACAGTGCGTAGGGCGCAGGATGCACTGCAGAGGACGGCGAAGTACAGTATCCGGGCACAGCCGTCGAAGTGTGCTATTGGTGGGCTGCAGGAGTTCTTGCGGGTGGATCGTGCTGCGGCAAGGCCTACCGGGTCTCAGTACCTGACTAGGTCTATAGCCACAGCTGTACACGCTCGTGCTGAAGCGAACGAGCCAGATGAGGCGCTTACCTTGCTCAAGGCGCACTACACGCGCCTTGACGAGATGAAGGCCAGGGGGGCGGATGAAGGGATCGTTGAGAAGCTGCGTTCTATGGTGAAAGAACGTATAGCCAGGGTATTCGCCCTAGAGGACGGTGTCCTTGAAGCGGTGGAGACAACACACTTAGTGCACGGGGGGCTGTGCGAGGATAGCGGTGTTGAGCCGAAGTATAGCGTGGTTAGTAGGTCGTCCTCTGAGATCGACGAGGAGACCGAGAGGTCTAAGACACTGCCGGGTGTCAGGGACTATGCTAGCTACCTGTGTAGTATGATGGGCGTCAGCGGCGCTGTTGAGGCGATGGCAGTGGATCGGATAGCTAGGGCTACAGCCAGGTTGGTAACATACACGCGCGTCGCGGTATACCTCCAGCCAGTTGAAGACAAGCTAAGAGCAAAAAACCAGCAATTTTTACACAAGGTTGCGAAGGGTGATTACGCAGTAGCGGCTTACAGCGGTAAGGCGCGCCTCGCCGGGCTACCGGTGGGCCGAGGAGCGATGGACAAGATGCAAGGAACGACACTTGCTAGAATCATGTCATCCCCCGACCCGTTGAGGACTATGGCAGTCATCTTCTGAAACAAGAAAAAAA

Color code: satar codon; sotp codon; slippery motif,

>Thecaphora_thlaspeos_totivirus_2

ACGACACTACCTTACTTTTCTCAGCTCGGGCGACACACACACCAACGTCAAAGCTGATAGCTTCTGGGTACGACACCAGATTAAAAACGTGGCAGCTGAAACAATCTGAAAAGCGACATAATGCAAACCAGCTTTTTAGAACAAAAACTCAACCTGACCTTCGGGCAGGAATACCAGCCAAAATTTGCCAACGGCAGGTTTACGCTGGTAAACAAGACCAACGCCATCGCCAAGTATGATGGCGTCGACACACAGGCCCTTCTTGAGCTAACGACCGACTTCATCGGCGTAGGCAGGAAGAAGCGCGTTGAGTTCGACGCTGCATACTCAGACTATTCTGGGATCAACGAGATCTACATCTCGGACAACGGTTCCTACATGGCCGATGCTGCGGTTGCCGACTTTGCCAAAAACCCTGCAGGGAAGAGGCTCACAACGGACAAGACTGACACCGTAATTCGAAGGTTGTCATTCGAAGACTCCCACCACTCCTTCATCTACAACATGCTGATCACATGGCTTAAAGCGTGGATGTACAAGAACAGTGGGGGTAAGAATAACAAATTCGAAGTGAAGTCTTCTCCGTACTCAGACTCGCACCGTACTGTACCGCTTGACCAACAAGCTGCAGATTACACCTACGAGATTGAGCTCGGTGTGCCAGCGCCCGGCACTGCAAGTGGCAACCCTGCCTCTCGTACACACGAGGGGTACTGGGAGTACAACAAGGTGTTCTACGTCAACAACACCACGAACGAGCAGCTTAATTTTTATTATGCTCACGTTCACGGTAGGACTGACGTGTCCGCCTTGAACTTCGATATCCCGCTACCGGGCATCGAAGGCTCTTGGTGTTTTGAGTGTGTCTCATCAAACATGGGGCACCTCGAACCAGAATCGGTTGACTACACCAACGCCGACCGTATTTGGGCCTGGATGATGGACTACGTCATCCTTAACAGGCTGCAATCCCAGTTTGCAGCCTGTTTCGAAACGTTGGGAGCGGCAGCCTTCCAACCGGTGTGGTCATCCCAGGAGGGATGTATGTGGCAGGAAGCGGAACTGCGTTTTTCGTTTGCCAAACTCTCGCCCACGAGAGGTTTGGTGCGTACGGCATTGGAGGGGACGCCATACCAGCTCAGCAACCTAGCAGCTGAGTTTATGTCTGAAGAGGCTCAATACTACCGGCGATTCATCGCAACTGCGTCGATCTTAAACTACTATATGTGGTACGGGATTTACGCGATGTTGGTCAATGAATGTGTCGGGCACGATACATGGCGCGACTGTCTGCACGGGATGACTGAGCAGACCGCAGTCCTGTCTAGACCTGAGATGCGTGCTTACGTCATCTCAGCGATCACTGGGCAGGAAATGGTTACGTTGATGACACCTGGCGCTAATCTCACGATTGACGTCAGTAAGGTCCTTGAGGTCCGCCGTATTGCCGGAGTTATTGTCAATGATAGCTCACCGGCAGGCGATATAATCATTGACCGACTTTATGCTCCTGTGTCGGGTTCTCTAGTACTCGGCACTTTCGGAGGTGAACTAGACGCAGTCACTCACCTGACTGCGATGCAACGATTTGCTCCTGTCGACGACCAAAGGTTGTACATGGATGATACAGACGCGCTTAGATACGCAAATCTGTATAGGCTGTTCGGGCACGATGTTAAGTTGCGCACCATGGACGGCCTACATGTTAAACCATGGGGCAACGTCAAGGAGGCTGTAATCGAACGAGGTTCGGTTATAGTCCCTGACGGTGCGGTCAGACTTTACCGTGCTGTGGACAGTGATGTGCGAAGTGGTAAACACTACCCTTTGCCACACATCCTAAACATGCTAAGGTCTGAGGTCACGCTTACAATCATGCGGCCGACTCTTGGTTTCTGCGAATACGGGACCAAGAAGACGCCGCTTGTAGTAAGGCAGGAAACTATGCGCAGGAAGAAAATTCCTGTTATGAAAGTTCGCTCTCAGTATGTCTCCGCGCCGCAACACCTAAAAGCGTTGCATAGGGCGCCGGCAGCCATCGAGCCGGGTTTTCAGGGGCCCGCAGCCAGCACGGTCGCACGAGCCCCAACTACGGAAGTCCACGACGGAGGGCTGCCGGACGAACCGCCTGCGATCACAACAGGCGGGGAATCTGTCTAGCATGCGCTCCTAGTGCGAAGACATTCGTAGGTGCCGGTGTATCGCCGAGGTCACTGTATCTGACAAGACCTGGGTGTATCACTGACGCCGTAGACTGTTCTGTCGTAGTGCATAGTGATGCTCCTTCACCTGATAACTTTCAGGTGTCGCGTGAGTTTGTCGACTTTGAAGTTTCGAACCTACCGGCCTACCTTAACGCACGATTTGACGCGGTTTCAATCGAAAACGCGACTTATTACTTCGTGACAACGTTGAAGGGTGAGTTCGCTAGGGGACACGTATCGTATTCGGTTGCCGGAAGCGTGATGAGGTGTTGGTCTATGCCACTGCCTGGTCACGGCGTCACTGCTGTGTATGCAGCAGTAGACCAGACCATTTTGCCCATGTTCACCTCAATTTTAGTAGCAGCAAATGCAGCGTTTGCAGGACACTACTACTATGACCTAGGTCTACCTGCGAGCATACAGTCTGTGTTTAACGTCACTGGTCGAGGACACTACCAGTACAAGACTTCTCTTGAGTCACAAATCAAGAACGCCTTCAGTGAAAAGGTCAGCGCAGAACACCACGAACACTATCGTGCCGGTGAGATGTATGACTTGATTAGCTCAAAAGGAAAGGCTCGAGCGAAAGCAGTGGGCCGATTAGGATTCCAATGCCATCATGCTATGTTGGCGGGCATCTTACTTTGGTGTGAAGTGCTGCCGGATGAACTTTGGACACAGGTCGTTTGCAGCAACTTGTTCGACGCTAATTCCGTCAAAGACTTCGGAAAGATTGGAAAGGCGATTTCTGTTGCCGCTAAGCAGCTACAGAATTTAACTGAGATTGACCTGAGGCCCATATTTGAGGTAGACGTGCTCGTCAACAGGATCACAATGAATGTTGACTGGGATGTTGAGCGTGAGAATAGGACTTCGCCCGTTTTGGCAGCTTTCACAGAAGAGGAGGTTTACGAAGCGGCAAGGCGTATCCTCTCGACTGAGGACAGCACTGTAGTTAGGCCTCGTGGTGTGAAGTGGGAAAACTTCTGGGCTAACAGGTGGCAGTGGTCGGCAGCCGGTTCGATATATACGAACCATCGTGCGGACGAACAGTTCTTACTCAAGGAACGTGCGTTTAAGAACAAATTCGTTACGCTCAACCTCTACCCTGACGTACCGTTAAGCCATTTTACGTCGCGCACACCCAAACTCGACGCACGAGCATCAATCAAATATGAATGGGCTAAAGTGCGTGCCATCTACGGGTGTGACCTCACTTCGTACGTGATGTCTACGTTCGCGTTCACCGGGTGTGAGGACAGCTTACCGAGCTACTTTCCAGTAGGCGAAAAGGCTAACGCCAAGTACGTTGGGGCTCGCGTCGCTGCAATGTTGGACTCATCATTGCCGTTCACGCTAGACTTCGAAGACTTCAACAGTCAACATTCGATTCCAGCGATGAAGGCTGTGCTTAATGCTTGGCTTGATGTGAATTCTGACAGACTTAGCCAAGACCAACTCGATGCGGGTGCTTGGACGGTACGATCTCTCGACATGGCTAAGGTACATAACAATTTGCCAGGTCACCAAGGGGAGTACAGAGCGAAGGGAACTTTGTTTAGCGGATGGAGACTGACTAGTTTTGTCAACACCGTACTGAACAAGGTCTATTCGGACAAACTACTTGCCAAGGGTGGTAGGATCGGCTCGCTTCACAACGGCGATGACGTGCTAGTTTCAACGCGTAACTTCAAGACTGCAGTCACAGCTCTCAGGAACGCAGCGAAGTTCAACGTACGATTGCAGCCCACTAAGACGGCATTCGGATCCATTGCGGAATTCTTACGAGTAGACCACGCTTCTACTGACGGGGGGCAATATGTCACAAGGTCTATAGCGACATTAGTGCACAGCAGAATCGAGTCTGGTCCTGCTTCTGATGTCAGGACCGCGCTAGAGAGTTATGAGGCAAGGTTTTCAGACTTTTTGGCTCGAACAGGCAGGACCGACATGGTGGTGAGGCTAAGAGAAAAGGCTTTCGGCAGGATGTCAGAGGTCTTCGAAGTATCTCCAAAGATGTTGTCGGAGGTTAAGCAAACGCATAAAGTGTGTGGAGGATTGGCTGAAGACAAAGCGGCATCAATTGAAAAGACTTTTAATGCCGTTGACGTGGGTACAGTGAACATTGACCCAGTCGAACTCAGGCGAGTGATGAAGGCGCCTGGAATCGCAGCGTACGCGAGACAGCTCGTTAAGCTCTTCCATCTGGATGATCAGTTGGCGGCGTTTAGCAGAAGACTTGCTGACGCGACTGTAAATGCTTGTGCAGTCAAAAGGAAAAGTCTAGTTTCCTCAAAAACGGACAATGTAGAAAGGCATACGGTATATAAAGCCCTCTACAAAGTGCATTCATACATCAACAAAAGTGGGACGTATGGGAAGGCAAAGATGGTTGGAATAGCTGTAGATGTATTAAATCTGAGATCAGATTGCAGTACATTGTGCACTATCCTAGCCAGATCAGCGGACCCTGTGACGGCCTTGCAAGTGTTAGTATAAATGCCAAAATATGGACGAC

Color code: satar codon; sotp codon; slippery motif,

>Thecaphora_thlaspeos_totivirus_3

CGAACATATCTACTTTTCATTGGCTTCTGGGTCCGACACCAGTATAAATACGTGGCAGCCTAACAAAACACTGTTCAACCCGAAATCTGTACGCAAGCAGCCAAAATGCATACCGATTTTATTAAAACACTTTTCAACACGCTCCTCGGGAGCAGAACATCACCGAGCTTTGCGAACGGCTGTTTCAGCCTTTGCAATGCTACCACAACGAGAGCGGTACATAAAGGTACCCATTACGCGTCGCAGCTTAAGCTGTCGACTGGTTTTACGGCCGCCGGTTTTCAGCCGACGGTCACGGACGTATACGGCGAAACGGACATCACGGGCATAAACAAGAGGTTTATTTCGGACGAAGGCACGTACAATCACTACGCTGCCGTTGCCGAATTCCAAAAGAATGTGCCCGGCTACAGGCTAACCGCGGTCGAACTGGAGGCGATGGTGGTATCATCGCTCATGTGCGACGCACAAGAAGGTTATTTGTTGAACCTGCTGATTAGCTGGTTCATTGCATTACTGGCTCAAGACTCAAAGACCAAGAATGATACTCTGGTCGTAGAGAAGTACCAGTACACTGACGCACACGTATCGTTTGGAGCGCCGTCGCGCAAGGGCCGGTCGCACACTGAGTACAAGCTTGGTTGTCCTGTCTCCGAGTACGACTTGAATGCCAACGTCAAACCCAGAGTCCGATCGAACTACTGGGACACACCTTACGTCGCACGGATCCCGGGAACGAGCAAGCAGCAAGTGTCTTTTTACGTGGCGCACTTGCATGGACGAACGCCAAGGTCGGAAATCAATGTCGACATTCCGTTGCCGGCCGTGGATTTTGACCAGCTGTCGGTCGAACTGATGGGCAACGTGTCCTACACGTTCGACTACAATAGTGTACCTTGGTCTAAACCTGAACAGTTGTGGACCTGGATTCGCGACTATGTCGTTCTTAACAGGTTAGAGACTGCGTTCGCCGCAGCGTTTGACCTGCTAGGCAGCATAGCATTCCAGCCGGTACCATCATTCCAGGAAAGTGCAGTATGGACACCTGCGGTACTTGAAATCAAGGTAGCACACTTCACGCCGACCAGAGCGAAGGTGCCCGCTAATCTTGAGGGCGAACCGTATCGCATGGATAACAACGCTGTCGACTTTGGAAAGCGCGAGAGTAGCAATGCTCGTGCGTTCATGATGACGTCGGCGGTGATGAACTACGCAATGTGGATGTCACTGTATGCAATGCTGGACAATGCGTCTGTGGCGTATCGCGACTGGAAGAGCACGCTCACTGTCGGTGAGCACGGATTGGAATGCACGCGCGGCATAGACGTGCGGGCGGACCTTGTCAGTGCTCTGTTTGGTAAGGAGGTCTTTACGGTTGCGACGCCCAACGCGCACGTGATACTGCACCCTGAGAACATAGTTCATAAGGGCAAGTATCAGTTCCCGTATGTCATAGGAAAGGGGTATCCGAGTCACTTTAAGGCTGACAGGCTCTACCCCTACGTATCTGGATCCCTGTTGTTGGGCACGGTGTCATCGCCGATGCGAGCTGTGCAACACTTGCATAGTAGCGCGACGGTGAAATTCGAGAACGATGCGACACTATCGAAGGTGGATTCGATTAAGCTCGCAACGTTCTATCGTCTGTTCGGGCACGACGTGAGCTTGCAACACAAGCACACTGGCGAGCAGTACCGCACGTACGCGCCACGGCTTGAGGGTGTGATATCACCTGAGCCGATGCTCAACGACGTGATGGAGTACGATCGTGTACTCGCGTACGAATCGAGCAGACGTGACGGCCGGAGTGAGAACTTGCCGTCGCTAGGGTCCGTCGCTTTCGGCGGAGAAATTGAGTTTCTGGTCGAGAAGCCGGTGCTGTCGCAGCTACGCTTCTTGTCTGTAAACGAGCGGAGTACCCCGTACTTCGTCACCGACGCGAAGCCCGCGCCGGTTGTTTTTAAGGTTGCAGGGCGCAGAGTATACACTCAAGCCGTGCTACCTCACGTCGACACAACGCGTATCAGGCCGGATTTTCAATTGGCCATAACGCAACCAACCCCTGCGGTGCCCGTAGTTTCGAAAGAAGCCACGGCTGTCCCGCAAACTGCGGAAACGGAGGCGGCCGATTTTGTCGAGCCTGCCGAATCGAGTGTCTGACACCACCGGCCGTGATCGGCGGTCGCGGCTTCCGCATGAAATTCAGTGAAATCGACAATGACTACGACCCCGTCGCCAGTTTCGTAACTGTTGGGGATGGCGATCGAGTGAGCGACAACTCGATCATTCTGGGGTGGTCGCGTGCCAAGCACGTGCCCACCATGATTAGCGTCCGGAACGGCAAAATTGCTCTTGCAGACTTTGCGGACGCTACAGCTGTATACGTATGCAGCAAAAAATCTCGGATGTGCGGGGTGTTCGAGATGGTGGTAAATGTACGCGGGACGGCAGTGTCAGTAACTGCTGTAGGAGACGGCAGCGTAGCGATGTGCTACGCGCATGTGGACCAACAGATGATGACGACAAACAGCACGCTGTTGGGTAGGATGACAAGGCACTTTAGCGGGCTCTATGCGTCGGTAAACTGGCTGGGGCGACTGAGTGAGAGCATGATATTCCGGAGGCGCGAATATGCCGGTGCGGAGGCAAAGACACTCATTAACATGCTTGATCAACCAAGCGCGAAAGTCACAGCGCACCATCACATTCATTTCACTGCTAGGGAGGTATTATCATGCGCCACTGCTGAGGAGGCTGCATGGTTACACGGAAATATAAAGTTGCCTAGCGATGCTACGCAAAGTTGCGTCGCCGGCGTTTTGGTGTGGCTCTTGGCTATGTCGCCCGATATGCGTGAGTGGATAGCTGCGAGTAATGTGTTGCAGGCGCCGAGTGTGAAGGAGTTTGCAAGGGAAGCTAAGGTGATGTCTGTCGAAGCCAAGTCGTTGCAGAATTTAGTCGATAGCGACCTGCGCGAGATGTTCGAACTGGATGTTCTGGTCAACCGGGGTATCGGGGAGGTCGATTGGTCAGAGGAGAGGAGGCATCGCGAGAAGCCTGATGTGGCGCGTCTCGACCCCGAGGCTGTGAGGCATTCGGCGCGCCAAGTGTTCGCCGATGCTAGAGCGATGGGACGCCGGCCGGTCAGTATGGACTGGAAGAAGTATTGGTCGTCTCGTTGGCAATGGTCTGCAGCCGGGAGCACGCATTCACAGTATGAGGAGGACATGTCGTATGTCGAACGTTCTGACATGCGGCTTAAAAATAAGTTCATCACGCTGTCAAAGATGCCAGCGTTCCAGTACGGACACTTCGCGGCTCGTAGGCCTGCACTACACGCTTGGACATCGTCAAAGTACGAGTGGGGCAAGCAGAGGGCAATCTATGGCACCGACATCACTAGCTACGTGATGTCGCACTTCGCGTTCTACAATTGTGAGAACGTGCTGCCAAGTCAGTTTCCGGTGGGCCGCGACGCTAACGACCGTAACGTCGCGAGCCGAGTGTCAGGTGTCCTACGAGATAGGTTACCTCTGTGCATAGACTTTGAAGATTTCAACAGTCAGCATTCGGTGCCTTTGATGCAGGCGGTCGTAGATGCATACGTGGACGTGTTCTCTGAGGACTTGAGCACAGAGCAGGTGATTGCTGCCCAGTGGACTAGCAAGTCGCTGGAAAACATGGTCGTGCATGACAACATGGGGCTGCGCACCACGTATTCGGCGAAGGGTACGCTCCTGTCGGGGTGGCGGCTGACTACTCTGGTCAACAGTGTGTTGAACTATGTGTATACTCGGGCCATCATGAAGGAATGGGGCTCGAAAGGTAGTTCAGTGCACAATGGCGATGACGTACTTGTAGGTTGCACCAACTTCGCGTCAGCGCAAGCGGCAGTGCGGAACGCACGTAAGCTGGGTATACGCCTGCAGGCATCCAAATGCGCGTATGGTGGTATAGCAGAGTTCTTGCGAATCGATCACAGGCGGGGATCGAAAGGCCAGTATCTAGCGCGTGCAGCTGCAACATTGGTGCATTCTCGTATTGAATCGCAAATGACAGTTGACGCAAGAGATGTAGCAAAGTCGATGGAGACGAGGCTGGAGGACCTGTTCCACAGAGGAGCAGATGCTTCGTTGATCGCGGCGTTACGCGACGTGTACTATAAGAGGCAGGCTGTGGTTTGTGAATGCGCAGTGGAGGACTTCTATCTTATGAAGAGGACACATGCGGTATGTGGAGGGCTGTCGAAAGAAGCTGATTCAGGAGTTGACTTCCTGGTTGAGCCGGGAGCGCGAAAAGTCAAGGAAGTTGAGCTGCCTCAAATGCCAGCGATTGCGGAGTACGCGGTCGTGGTGAAAGATGCTCTGCAAATAGATGTGCCATATAGTATGGTGGTTTCACGTCTAAAACACGCAACTAGCGAAGCGGTGACTGAGAAAAGTCGCACGATGAAAATCGTGCCTTGCTCGGATCCGTGGTACAAGCGTGTGAAACCAATTTACAAAGCACATCGCGGCAAACTGAGTGTTGCCGAGTACGGAAAAGCGGCTCTAGTGGGGCTAGCTTATGATGTACTGAACGCGTCCGAGGGTGATAGTACTCTCAAAGCAATTTTACGGTGCTCAAAGAGGCCGATACAGTTGCTTGAGATGCTCATCTAAGGTGACGCAAAAAG

Color code: satar codon; sotp codon; slippery motif,

>Thecaphora_thlaspeos_totivirus_4

CCCTTATAATCCCCAATGTCTTTACTACTCAAGCTCGCGCACCTTCAGAACATCTCGACGCACACGCCGACTGTGCCGGGTGGTAGGTTCTCTATCACTTCGGAAACAGCAGCGGTGCTCGACCACCAAAGCACAAAATACGTAACCGACCTCTCGCTCAGGTCAGATTTTTCATTCGCAAACAGAGGCAAAAAAGTCTCGTTTGTTCAGAATTATTCCAACTGGGCAGGGGTCAACAAAAAATATATTGATGACACAGGTGATTTCAACATGGCCGTAGCCATCGATGATTTCGCCAAGGTAGCGGCAGAGAAGAGGGTAATTCGCGGCGATGTAGTCCGACAGTTAGAACAGTGGCCGAGGCGTGACTCACATGAGGCATTTTTGTATAACATGCTAATCACGTGGGGGCATGCTATGCTCTACAGGCTAACAAAAGGAGAAGAGGGAGTCTACAAAGTGGTCACCAGACCGTATTCGGACTCTCACGTCATCGTCGCAGACCACCTCGGCTACAAGGAAGAAGTAGCACATGAAATCGAGATCGGAGTACCTGCTATCAACGCGGGAACTCCGAATTGGGTACTACGCGAAGAAGAGAGGTTTTGGGACCGCAGATACGTTTTGCGGTATTCCAACAAATCTAGCGCCCAGGAGGCGTTTTATCTTTCGCATGTCTTGGGCCGTGATCAGGTGTCGATACTTAACTTCGATGTCGCGATCCCGTCTCTAGATACAAGTCAGTTGATGCTAGACCCGATAGGTACGGCTGTCTTGGGGGGTGGAATGGTTGCAAATCTGGATGGAGAAGGGTTTGAAGACCCTGACAGAGTGTGGTCTTGGATCATGGACTACGTAGCTGTCAATCGACTGCACATGCAATTTGCGTCGGTTTACGAAACCTTCTCTGCGATGTGTGCGCAGCCAGGCTACAGCTCTGCTGAGGCCTGCGCTTGGCAACAGGCATCGCTATCACTCAGGTTGGCGACCTTCTCTCCAATGCGTGGTGTTATACGCACCAACTTTGAAGGGGAAGAATTTTGCCCCTACCCGGACACCCTCTCTATGCTCTTGCGAGAGGGGAGGGTACCGGACCAGTTCGTCCAAGCGTCCGCTTTGATGAACTACTACCACCTTTACGGCACCTACGCGATCTTACACAACGAAGCGCGTGCTCGTGACCAGTGGAGGACGGTGTTTGAGAGCGCCGAAGACGCTCTCTACAACCTCGACACACCCGTAGCTCGTGCCACTGCTATATCAGCGGTCATCGGACAGGAGGTGTCGACAGCCATGACCGACGGATGCTACATGACGCATGACTACTCAGACATGCGACGTGTCCACCAGATCACTGATGTACAACCGGTCGATGGCTCAGTAACTAGGTTCTCTGTTGATGCCCTTTACGCGCCAGTTGACGGCGCGTTGCTCCTGGGCGTTGCAGCTGGCGGACTCGAGACGACCGACCATCTCAGGTCGGTTCAGACTTTCGAGCACCACCCGTTTGACGGGGACCCACTTACCGCAACGCACGCCCATCAGGTGGCGGCGGTCTTCAGACTATACGGCTATGACACCGTATGGTCTTCGCGTTTCTCGCGAGAGACCATAGTCACTTGGGCCAATGTTGCTAGTGTCATCGTCGACCCGGCGGTACTCTTCTTCGACCCAATGAAAGAAGAGTACTACCATCCGGTAGATGCTGAGCGACGAATGGGCAGGTCAAGGACCCTGCCAAATATCGCGAACTTCCTTGAAGATGGGGCGACTCTGCAGATACAGAAGCCTCGCTACAAGGTCACTGAGCACGGAGATAGGAAGATGCTCTCCAAACCCTACACCAGGGTGAATCGAGGCAGAAAACCTATAAACGTCCTCATTTCCGCGCCTGTGGCATACACTTCGGCGCGGTATGTTGCCAAGCCCGTTAGACACAAGCCGGTGCAGGATTTTCGAGGTCGCGATCTCCCTATAACCCCGGCCAACCCCGAAGGACAGCGAACAACTACTCGCGTCGACGAGGCAATAACGGACACTGCGACCATGGCACAGACTTTGACCGATGCACCAGGTGCAGGATAGAAGCCTTGCCGCAACATGACAGCTGGACAGTGAGACAAACTCGGACACTTAGAGCTGGCGGGCACGTTGAACGCTCCGACTTCGGAGTGTTCTACTACCCGCCGAACCTGGAAACGAGTGGAGGCGTGGCTTTCAAGGGGCACCTGAACACGCTGGGCAAAGGAAGAGTGCCTACTGCTATTCAAATCGTCGACAAACTGACGGTTAAAGCAGTTTCGTATAAGGACGCAGATTATGTCCTTATCGACAAACTTGACCAAAAGTTTGACGATATGGAGGTTCTGTATTCGTACTTCGGGTGTTCGGTCCATTGCAAGATGTTGGCCGGCACCGACGGTACGTATGTATATTATAAAGTCGACCAAGAGGTGATGCCGGGCAGCACCGACTTGGCCGCCGTGATGTCCAGGCATTTCTCTGGAGATTACGATTTCTATTACAACTCGCCTAGCGCGAGCGAAAACCTCACACTCCCTTTCTTAGGCAGGACTCAGAACTCTAAGCGCTGGCGGAGACTAACAGACTTACCGACCAATAAAATAACGTCAGCGCACCACACTCATTTTAGGGCCCAGGAGATTTGGAATGTGCTCGACAACGAGCAAAGACGGAGGGCGACAGAAGTGTTCAGATACCACGGCGACGACGTCACCGACTCGTTCGTCGCTGGCTCGATGCTGTGGTTAGCTTCGATGGAGGACAGGCTATACGAGCATGTCTGCGCCATAAACTTGTTCTCAGTCTCGAGCATAGCGGAGTATGCTGCGCTGGCGAAGAAGCTCAGTGTACGCGCAAAGTCGTACCAGAACATATTGCATGACGACCTCAGGCAGTTGTTTGAAGCCGACGTGCTCGTCAACAGAGGGGTCGGAGAGGTTGATTGGAAACAAGAAAAAGAAAACAGAGTGCGACCTCAGCTGGCCGACGTCAGCTACGATGAGGTGTATTCGATCGCTTACAACATGTTCGGAGACGCGCTTGAACAAGGAGCGCGGCCACGGTACCTCAAGTGGGAGGACTTCTGGCAGACGCGCTGGCAGTGGTCGGCATCTGGGAGCATACACTCCCAGTATCCGGAAGATCTAGTGGGCTTACCAACCGACAGGCACTTGAGGAACAAGTTCATCAAGTTGGCTCAGATAGGGCCACAGCCTATCAGTAGCTTCATAGACCGTACGCCGGAACTACAAGCTTGGTCGTCTACGAAGTACGAGTGGGCTAAAATGCGCGCGATTTACGGCACTGACCTCACGAGTTACATCCTTGCGCAGTTCGCTTTCTTCAACTGTGAAGAAACGCTTCCTGCGCACTTCCCGGTCGGAAGCAAGGCCAGGGACTCGTATGTAGCTGCCCGTGTCTCAGCCGTGTTGCACAACGCTACGCCGTTCTGTCTGGATTTCGAGGACTTCAACAGTCAACATAGCGTAGAGGCTATGCAGGCTGTGCTCGACGCTTGGCTTGCGAAGTGCGGACCTTCGCTCGCCCCCGAACAGAGGAGTGCGGTTGCATGGACACGTGCCAGCCTCGCCAACATGCGTGTTAATGACCTGTCCGGTCTCAAGCAAAGTTACAAAGCAAACGGGACTTTGCTGAGCGGATGGAGGCTTACTACGTTTATGAACAGCGTGTTAAACTACGTCTACACCACGAAGTTGATTGGGCAGGGCGGAACACAGTACAGGAGCGTCCACAATGGCGACGATGTTTTGATAGGAATGACGAACTATTCCCTCACGACACAGGCGCTACGCAACGCAGTCAAGTACAAGATCAGGATGCAGCCTGCAAAGAGCGCGATCGGCGGGATTGCAGAGTTCCTCAGGGTGGACCATATCCGTGGGCAGGACGGACAGTACCTCACAAGGAGTATCGCGACCCTCATCCATTCGAGGATCGAGTCTAACCTCCCGGTCAGCTTGATAGATGTCGCGAACTCGATGGAAGAACGATTTTCGGAGTACGTCACGAGGGGAGGCAACTACAGAGTGGTCGCATCACTCCGTGACAGGTACTACCACAGGATGTCAGAGATATATCACTCTGATGTCGAGACCATGTACATTATCAAGATGTCGCACAGGTTAGTTGGCGGAATATCAACGATGCCTGACGCGATGATGAAGTACAAGCTGACGGCAGACAACACGATGAGCAAAGTTGATTTGCCGCGTAGGCTACCTGGAGTGGTAGACTACGCATTTGCGCTCCATAGGCAACTGGAACTACCAGGCTCGCCGATGGAGTTAGTAGATAAGGTGATGACAGCCACCAGAAGAGCTGTCCAAATGGTAAGGTCTAAAGTAACACCCGTAGGCAAGGTTGACGCACATCGTTGCGCAATATTACGTGGAGTGTACGGAGCTTACCGCGACCTGAGGACAAATACTCAGTTCGGAAAAGCCAAGATGCTGGGCATAGACGTCGACCTACTTAAAGGAGGTGGACTGCTCAGCAGGCTAGTGACAATGTTACGTAGTTGTCACGACAAACAGGCCTATTTAGCCGCAGTGTTATAAAGACGCTGGAATATAATAAAGGTGA

Color code: satar codon; sotp codon; slippery motif,

>Thecaphora_thlaspeos_totivirus_5

ATTAATCCCCATGGCTCAGAACGATTTTCTCCAGACTTTTGTCAATGATTTCTTCGCGAGACAACTCTCGCCAAAAGTGCAGCAAGGGCAGTTCTCTATCGTGAACAGGACCCATGCAAATGTGTCATTTGAAGGTGGAACCACCGTGTCGGACCTGAAACTCCGTGCGGATTTCCAGATAGTCGGGAAGAGGAAGCGGCTCGTCCTCGACGAACTCCATACTGATTGGGCAGGCTACAACAATGCCCTGATCAGTGCGGACGGAGTATATCAGCCGAGCGTAGCGCTGACCGAATTCATGAAAACGGATTTCGGCAAGCGTGTGATGCCCACTGATATAGAGGCCAAAGTACGAGAGTTGCCCCAACGTGATAGCCATGTGTCCTACCTGTACAACATGCTTATCTCTTGGATGAAGGCAACCCTCTACAAAATGAACGACGGCAAAGATGACGTCCTGCAGGTCAGGGAGTCACCTTACCGTTCGTCACACGTGACAGTCGGCTTGAGCAACACGACGGACATCGTACACGAGTACGACCTCGAGGAACCGGCCGACATCAACTCCGCACCACAGACAAGGCGAGATAAAGACAACTACTTCTCGACCCCGTACGTGCTGCACTACAACGCGTCGAATAAGTACCAAGAGTACTTCTATCTCGTCCATGTGCCAACGAGGAAAAGCACAACCAACCTGAACTTCGACATTGCAGTCCCTGGACTGGGTGCGAAGGCGGTGTTGCTCGACCCCGTTGACCCGTGGGCGGATTACGTCGACGTTGAGTGTGGCGGTGAGATAGATTGGGGTGAACCCGACATCATGTGGCAGTGGATCATTGACTACGTGTCAATGAACAGGTTGGAACAGCAGATGGCGAGCGCTCTGGAAGTGCTAGGGTCTGCATCGTTCCACCCTATCTGGTCGACAGCTGAAGCATGCGCTTGGCAAGCGGCAGAACTAAGTTTCGTACTAGGAACGTTCTCCCCGACCAGGGCCCGACTAAGGACTAACCTCGAAGGAGGTAGTTACACCACTACTCAAGAAGGAGTGGATTATTATATGTCGAGCAAAGTTCAGCCAATGCGGTTCTTGATTACCGCAACAGTGCTGAACTACTATATGTGGTACGGCCTGTATGCCGTTTTACACAATGAAGGACGCACGCGCACAGCTTGGCGTTCAGTCTTCGACAGCACCACAGATGAGCTGTCTTTACTGTACACGCCATATATGCGCGCGTTCTGCATTGGGGCGATTACGGGCCACGAAGTACCGACGGCCATGACGCAGGGTTGCGGGTTGCGAATCCGTACTGCGGACGTGGCAAAAATCAAGAGGATAACGAACACGGCGCCCTCATCTACCGGTGAGGAGGCAGCCGTTGTTATCAACGCGATCTACGCGCCTGTGACGGGCGCATTGATCATCGGATCGTTGGCAGGCGACCTTGACGTCGTCTCACACATGAAGTCTCAGTACAGGATAGACTTCCACGGTGTGGGTGACTCGCTGTATAGCTTTGAGGATATGCTCAGAGTTACTACGGCCTACAGGTTGTTTGGCCACGATGTGACACTCATGGACACGAGGACGGAGGAGCTCGTCCAGCCGTACGCGAACCCGGCTGAATGTATCCCGGAGCCGGCGTCCATCGAGTGGAACACCATGCGGACGAAGCCGCTGACTATCATGTACAGCGACCCGCGTGAAGGCCGTAATGTCGGAATGCCTGCCCTGTGTAGTGTAGGGCTGGGCGGTCACATCACGGTCACCATCGGCATACCGAGCTTGGAAATCACACAGTTCCAAGCTGCGAACCGGCCGATGGTAATGTACAACAGGACGAACTACAAGAAGAAGCCTGTTGAGATAAAAATCAAGGCAAAGTATCATACCGTCAATCAGAAGTTCAGGGTGACTAAGACGGCGAACCCTTTGCCGCAGGATTTTACCATGGAAAGGAGCAGCCTACCATCCACACGCCCAGAAGGGCCGAAGGTAGAGGAGAGGGCTCAGCACACTGTCCCTGCGGAACCGACATACGCAAGTGTCGCAGGTGCGGGATTAGAGAGTGGGCAAATGTCGTCCGCATTGGAGACGAAGCCGTCTTCAAGCGGAGGAGCGACAGGAGAACAGACAGGCGTGAGCTCAACGTCTGCAGGATAGAAAGAGCACCTGAGGGGAAGCGGGCTAATGCACTAAGGTGGACCCACATTAGCACAGCACCGTTATGCCTCAGGTTAACAGACGCGAACATCTACGGGGTCGAGTTCGCGCAAGCTACGCACGTGTTGGTCGACTGTGTAGCTAAAGGCCCCGAACGGGGCGAATTTGTACATGATCTGTTTGGGACAGCTGTGAGGGCGGAATACATTCCGTCGCCACAGGGTACCCTTGTATATATGAAACTAGATCAAGCGATTAATCCGATAGCAGGTGAAGTTACAGCGTTACTGGCAGCACACTTCGCAGGCTTCGAAGTGTACTACAACGCACCCACGATCCTTGACAACATTACCATCGGGATCGAGGCGACCAAAGAAAGAAAACAAAAAATTGTAAAATTAGAAAATCTACCAAAAACAGCTATTTCGGGCGAACATCACATTCACTACACAGCACAAGAGGTTTGGGATGCTCTTGATGTACAACAGCGGGCGCAAGCAAGCGCAGCACTGTCCTTACCGCCTGATAGTACCACTACGTTCATAGGGGGCGTCATGTTGTGGTTAGCACAACTTGAAAAGCCGCTCTTTGACTTAGTGAACAACACCACACTCCTTGACAGCAGGGACTTCAGAGAATTCGCGAAGATAGGAAAACAGATATCAGTACGGGCGAAGTCACTGCAGAACATGACAGCAGCAGACTTACGTAAGATATTTGAGATCGACGTGCTTGTAAATAGAGTAGCAGGAAGTGTGGACTGGGAAGGGGAAAAACAGAACCGCGTAGAGCCAAACACTGCGCAGGTTGACGTCGAGAAAGTTTACCAGATGGCGATAGGACTGTTCTCGACACACGACGCCAAGGCAGAGAAGCCGAGGAGGTTCCCGTGGCGCAAATTCTGGCGCTCGCGGTGGCAATGGAGTGCGTCAGGCGCTATACATAGTCAGTATCAAGATGATCTGCAGTATGTAGCGAAAGAAAGGGAGTTGAAGAACAAGTTCATCTCTCTAATCAGGATGCCGTCGGTGCAGTTGGAACACTTCACCGAGCGGAAGGCATCCATGGAAGCGTGGGCATCGACCAAATATGAATGGGCGAAGCAGAGAGCGATCTACGGGTGTGACCTGACCAGTTACGTCCTCTCGCACTTTGCATTCTTCAACATTGAGGACACTTTGCCAACACAATTCCCGGTCGGTAAGAAAGCACGGCCAAGCTATGTTTCTGCACAGATATCGGCGGTTCTTAGAGATAGGATGCCGTTCTGCGTAGATTTTGAAGACTTCAACAGTCAGCATAGCACGAAGGCGATGCAGGCAGTACTGCAGGCATGGCTGGACGTAAACGCGCGTAACCTACACGAAGATCAGGTGAAGGCAGCGCTCTGGACCATACAGTCGTTAGAGGAGGTATTCGTCAACGACAGTATCGGGACGGGCACGCGTTACAAGACCGACGGCACGTTAATGAGCGGCTGGCGGCTAACGACATTCGCGAACAGTGTACTCAACTACATCTACTCGCAAATGTTACTAGAGGGACACGAAGGTAAAGTTAGCAGTGTCCACAACGGCGATGACGTTCTGTTAGGAGTAAGTAGGCCTAGCGTGATGAGCGACATCGAGAAGAACGCGAAGAAGTACAACATACGACTTTCGCGGAGCAAGTGTGCATTTGGAGGCATTGCAGAATTCCTCCGTGTGGACCATAAGCGCGGGCAGTACGGACAGTACTTGACCCGAAACGTGGCTACGCTAATGCACGGCAGGATCGAGACAAAGCAAGCAATTTCTCTGAAAGACGCAGCTGAAGCAAATGAAGCACGTCTACAGGAAATGGTGGTGCGAGGAGGCGACCCGCTAGTAGCAGCCAGACTAAGGGAAGGCTACTACAAGCGGCTCGCGAAGATCTGGTCAAGTGACGTAGAAGCTATCTACGACATGAAAAGACTACATCGGGTGGTCGGAGGATTCAGTGAAGAGTCCTTCGCCCCAGTCGATAGGTTGGTTGAAACCGAAAAAACAACAAACACGGTAGAGCTTGAGGAAGCACTACCAGGGGTATACGATTATTCAGTGGCCCTAATAGAACAACTGAACCTTAAAGCAGGAGTAAAAACTGTTTACAAGCGTATTTACAACGCTACACTCGATGCGGTCCAACTCACGAGGCAGAAGACTACGGTCGTGCCGAATGACAGGATGCAGCAGTACGTGGTGTACCGAGGTATATACGGAGCGTATAAACAGGTTAACGACAGCGCCCTAGTAGGTAAGGCTTTACTTACAGGGTTCGTGGTTGATGTGCTTAGCAAAGCAGAAGGAGTGGCAGCTCTTTCTATGATGGTGAGCGCAGCCCCGAATCCGATGGAGTACTTACGCGTCGTCTTATAATAACCTTGAAAGCTCAAAAAGGCGG

Color code: satar codon; sotp codon; slippery motif,

>Thecaphora_thlaspeos_eimeriavirus_1

GTTTAGCGTCTATCTTTCTGTTCCTGTTAGTAGCCAGTAAGCTGAGTTTCGGACCTGAGAGCCGCGATTCTTTGGAAAGGGAGACCGGATGATACGGACGTTGATTTCACTCCTGGGTGTGCTTGCATTGGAAAGTCGGAGGGATTCCGGAGATGACCGTACCAATGCAGGAGACTAACAGTGTAACATCGGCTTCAGCAAGTGTTTCTAAAACCCACATGTTTTCACCTGTGGGTAGGGCACTTGCTGCAAGCGAGATTAACAACAATGGTCGCTTTCGTACGTACTCCAGTGTCGTGACGAGCGAGATTACCCTTCAGGGTAATAGAGACATCAGCAATAGGCTGAATGTCTATAAAATTGGTGCACGGTACACCAATACGGGCATGAATCCTCATGCCCCCTCCCCCCCGAGAGAGGACTCTGGCCGCAGAGTGATGCTCGGGGGGGCTGATGACGGTAAGGAAAAGAAGGATCGCAAAACCTTCGTGGACATACCCCCTACGCCTTTGGCAACAGGGAAGGTCCATCCGTCGGAGCGCAGATGGGCACAACCGATCCGGGTGCCTGTCGCAATTGCCGAAGACTTCGCTAAACAGGCGAAGAACTACAGCAATTTCAGTGGGGACTTTGCGACGGCTGATCTCTCCGGGCTCGTTTTTCAACTTGCTCGCGGGTTGGCCTTCTACACCATCAGTCCTACGTTCACCACGTATGACTTGAGTGCAGGCAACATCCCCCGCATAGTGGCAGTGAGCATGGTCTCAAACTCATTGAGTGCGAGCAATAACCATGTTTGGTGCCCAAGGCAGATTGACGACAAAACGTCGCCAAACGTTGTGGCAGCACTTGCTGCAGCTTGTGCTGGTTGTGGGTCGACATTGACTATGCCTATAGTCAATGTTGATATCAACAACAATGTGGTGGTCCCAACCGCGGCGGGTGAGGAGCTGGCCTTCGGCATCTTCCATGCGTTAAGGATCATCGGTTCCAACTATATGTTGGCTGGGGCGGGAAGTGTTTTTGCATATGCGTTAGTTAAAGGACTACACGCAGTACTCTCATTGGTGGGCATGACCGATGAGGGTGCATACATGCGCAGTGTGATGAGGTACAGTGCCTTTATACCTAGTTATGGAGGTATTAAAAGTGATGAGCGTCAGTGGCACGGCATGGCTGAACCAATCGTCGGCAGCAGGGGGTCGTGGACTGCCCTCGTCGACGGGATGTTGATGATGAGCGGAGGGTTGAGCGCACTGGCCGACCCTTGCAGAACGATTGGGGATAAGGTATATCCCTACACGGTGGCTGACTTTGGCGCTGAGACGATGGCTTGTGGCGGGAAGAGTGATGGCGACCAGCCCGGTGCAATTCGGATGGCTCGACGCCTCTTACAGGCGAGTGCGCAGTGGAGTGAAAACTACGTTCGCTTGCTGGGCAAGACCTTTGGGCTGATCGATGGGCGTAGCAGCCAGTTTGATCAGACAGAGGAGGCTGTAAAGCATTTACAGCAGTCATTTGCCACTGGTTGTGAGAAAGGCGATAGGCATCTGACGCACGGTAGAGTTGCTTCACCATTCTACTGGGTTGAGCCGACCGGGCTGTTTGACTGGATTGATTCCACTCTACCAGCCCAGAAACACGGTTACGGTGTGCTGGCAACACCGGACTTGCCAGGTGAGCTGCCACTGTTCCCGAAAGCGCAAGTGGTCGCTCGTAGCGGTGACCTTACCGAGGTCGAGTTTGAGTGGGTCACTGCGCGGCGCACTGGTGCGTTCCACCATTTCAGGAACCATAAGTTGGACGGTCTGGGTTACATTCGGCCGTTCGATTTCAACCCTGATGCGTGGACGAGTGTCGGCGGGGACGGGTCGTACATGTTCGCGAGAGCTGAACAGTACAAGGATGTCACTAATTACCTCTGGGGTAAAGGTGACAATGCATTCACGGCCCCCGGGGAGGCAATCTACTTGGGGGGCAAAGTAAGCGTTGTGCTGACGCACAGTAAGGCGTGTAATGCTTACATCATGGAGGAGACCCACATGGTCGACAGCTCCGAGGTTGACCAGCTCGTCAAGATACGGGTTAGCAAGTTGCACGCTGAGGGCATTCAGGGACTTGGTCCGATGCCAAAGGTGCAGTCGGCACTCTCGCTGGCAGCGGCTCAGCTGGCGGGAGCGACGAGGGCAATGGAGCGTCTGGGGCGTGGTGCGTTCGGCGTGCCTCGCAGACGCATGACCGGAGAGGAGCCGTTCTGTGACGTTGACAAAGCACCTCCTCGGGAGAGTAGTGCTGAACCAGTTCACGCAGCGGCTTCGTTCGTGCAGACGGTGATCATGCCAAGTATCCCGTCTGGAGGTGTGGCGCAGCAGTCGTTGGGCGAGATGAGCGGCAACACCGTGGGTGTTGTGCGTGAAGCGTGTCCCGAGGGTGCGCCGCGGAGCAGTAATGCCCAAAGGGGTGTTGCGGCTACGACAAGCCTCGTCCCTGAGGCAGCAGCAATTCCGGCTGCCCCTGGCGGTGATTCAACCGCTGAGGGCGGCGTCTGGAC**ATGA**GCGAGAAGCTCTCCAGGCGTCTGGAGGAGCTCGGGCGTCTGGGTAAGCAGCTCCTGCCCCTGGTGCGCCCTTACGTGAAAGGGTGGCCAGAGGATACATACGGTCAGTGGCGTGAGGTCGACAGTGTTCCCGGTACAGGGCTGAATGCAGAAAAGTTGCGAGTGGGGTTGTCGCTGTTGTTGTGCGATTACCCCTACCAAATGAAGATATCAAATACTGTGCTCTGGTGGCTTTTTGACAATTGTATAGAACCGCTCGAAGATATGGCTACCGTTGAGAAAAGGCTCGGCAAAAGAGCAAACGAAATGCGGGAGGGTCTGGCTGACAGTTACAAGCTTAAGATCTTCCCCTACAAAACAGATGTGGCTCGTGCAGAATGCAAGACAAATGTTTGGTTCAGTTCCGCTTTCGGTGAATTGTGTAGGCGTGACAGAGCTTCCGCGGAACTCTGCAGATACTACGCTCCTTACCTTGATGGGTTTAACGATAGCCAGGCGGTGAGTTGGTTGCTCTACAGCACCGCCTTGGAGCGTTTTACCTCTGATGCGTTTTTACGGGCTACTGCCTTGATGCTGGACATCAAGGGAGGCAAAGTCCTTACAAACTGCCTGAAAGGCTTGGGGATGAACGGCACAGCGGTGGGCGCAGTGTTGTGCGAAACCAATGCTTTACAGGGAAAAAACGTGGCTAGACTTGACTTGTCTGTTGAAGCTAGCCAGCGATGCAGTGAGAAGTGGGTCAAGGAGAGGATGTACCTGCCGGACATGGAGGCGTTACGTGTCGCTGTAAGGCGCGTGTTAATGGACGAGGTTGACATGTCAAAGTACGATCCTCCCACAGATAGTGAGTGGTGGTCAAGCCGTTGGTTGTGGTGTGTCAACGGGGCCCATTCCCGTGTCAGTGAGAAGAGGCTACACGGGGGAAGCGTGATCCCGAGCGAGATCGCAAAGCCGTATCGAAAGGTTTATGCTGAGTGCAAGGAGGCTTTCGATATCGACAATTGGTCTGGGCGAGTGGAGGCAAGCGCTAGTGAGAAAGCCGAGAACGGGAAGCGTCGCGCTATTTTTAGCTGTGACACAGACAGTTATTTGGCTTTCGAGCGGCTTCTCAAGCCAGTGGAGGACTGTTGGCGCGGGCATCGCGTCGTTTTGTCCCCTGGCAACTTTGGAGGATATGCGATGGCGAAGAAACTTCGGCGCTTCAGTGACATGGGTTGTACTAACATCATGGCTGATTTCGATGATTTCAACAGTGCGCATAGCCTCGAGAGTATGAAGTTGGTTTTTGAGGAGTTGACACAGCTGGTCGGGTATGATACTGCCCGTGCGGAAAAGTTGTGCAACAGTTTCTACCACACGTATGTGCGCGGGGGACAGTTCGAATGTGGTAGGGTTGTCGGGACTCTGATGTCTGGTCACAGGGCTACGACGTTTATCAACAGTGTGCTTAACAAGGCATACTTGTCTGTAGCAGTGGCAGGGTTTGATGACCTGATAAGCATGCACGTTGGCGACGACGTGTACGTTGCGGTGCGCGACCGCAAAATCGCTGCTAGGGTCGTAGACGCGCTACGGGATTCCCCACTGCGCATGAACCCGACAAAGCAGTCTGTAGGGGGTGTGACGGCCGAGTTCTTACGAATGGCTGTTTCTGGTGAAGGGGCATTTGGTTATGCGTCGCGTAGTATCGCGAGCTTGGTGTCAGGCAACTGGACCAATGAGCATAAACTGGACCCCGAGGAAGGACTGACTAGTTTAGTGAACAGCTGTTGGTCGTTGCAGAACAGGTGTATGAGTGATGGGCCCTGTCTGCTAGTCGTGCCCAGTGTCGTAGCGGTTTCTGGGCTAGACAAAAAAATCGTACAGGACTTAGTCTGTGGTTTAACTGCCTTGGGCGACGGTCCGGTACGTGGCGGTCGCAGCATTTACCATAAGGTAGTGTTGCGCCGTGAAATTGCGGAAGAACGGAGTGAGCAGTTGGCTGTGGGTGAGTTTCCACACCACGCAACGAGCTCGTACTTGCAGTCACACGTAAGTCCCGTGGAGATTGAAGGTATGCGTCTTGCTAAGGCAGACATCTCTCAGGTCATGGTAGACTCTAGCTATCGCAAGAGTCTCGCTGAGGACGATGGATGTACTATGCAGATGCATTGGAGCCTTGAATTCTTCACTCGGCAGTCACTGGCGGAGCGAGGCGAGAGGTTGAGTGACCTGATGGAGCTACGGCCGATTGAGGGTGCTTTGACAAAGTACCCTCTGCTCAATCTGGTTCGCCGGTCACTGCGGCGGCGGGATCTAACTAGGTTGATCCTGCTGGCCGGTGGCATGATCGGCAGTGATGTGGAGATGAGTGCCTGGGGAGCGAGCGGGCACGGCGTCATAATACACGGTAATTTGCCGCGTAGTGACGCAATCAAATATTCCAAGCGGACGTTGGCGACGTCGTTGGTCTCTGACTTAGATGTAAATTTCTAAAGTCATATAAATAGCCCTTGTTAGCAAGTAAAATTTGAACGAAGACACGAAAGTGTCGC

Color code: satar codon; sotp codon

>Thecaphora_thlaspeos_eimeriavirus_2

AAAAAGCCGAGGCTTACACCCCTGTGTTGCCTCGCCGCGTGACGCCGGAGAGATCTGGTTGGTTGACCGCGTAGTTTTTTATCACAACACTTAATACCACCGAGGGGATTGGGCCGCAGTCAAGGACTGCCATACGATTTCCCGCCTATCGAATATCTGTACAAGTAAGGGCTTATAGGAATCAAGGAGGTGCGTGTGGATCGAAGGAGGGTTCCGTTGATCGCGTGAAGCACGTAACCAAGTATTAAGCATCAACAGAATTATAAAATCTTGCCGCAGGCAAAACAACAAAGTACCATGTCCGACGAGGCAAAGAACGTTGCTTCGTCGTCGACAAGGACATCTTTGTTGACGGGGGTGGTGTGTGCGCCCATCGCTGGGGCGTACGGTGACAAGGTGTTCCGCCGTTACAAAGCGGATTTGACGTCGTCGGTGGTCATGCGTGGTACTGTGGACACGTGGACACGAGGCATCAAGTATGAGGTGGGGGCGCGCTATACGGATAAGGCGCCTTACTTCGCTGAGTCGGGGGAATCCATCACCCCCGACACTTCGGTTAATACGAACATGATGTTAGCCGCAGACTTTTCAGGTTTCGCAAAGAAGTACACTAACTTTAGCGGCACCTGGAACCTGATGGACCTTTCCGCGGTCGTAGAGAGGTTGGCGATCGCGGTTGCGGTGTCGTCCTGGTGTGAGGATGTGTCGACAGACGACTTGCGGGGAGGCATGCCCGTGAACGTAACTGCGCTCGGCACCCACTTTTCGCCGGTGTCAGCAAGCACTAGCCAAGTCTTCGTGCCGCGCCTAGCGGACGACGTTATATCACCGGACGTGTTGTCGGTGTTGGCTGCAGCTGTGAATGGCGCTGGTTCGGGCCTGGTGACGGACTTGTTGGAGGTGGACGTGCGTGACAATCGCGCAATCGTCCCGTTCCTGCGCGGGCGCGGTATGGCTGCTGCTTGTGTGGAGGCGCTCCGGATCATCGGGGCGAACTATTCCTTGAGCAACGCTGGAGCTGTCTTCTCGCTGGCGTTGGTGAAAGGTATCCACAAGGCCGTAAGCGTTGTAGGTCACACTGACGAAGGTGGCTACTTCAGGCGGGTGCTTAGGACTGGTACGTTCCGGGCACCTTACGGAGGTATCCACACGGAGCATCGTGACTACGTAGGGATACCTGCGATCGCAGGAGAATCGATGAGGGCTGCGGCAGGTTGGGTGGATTGTGTTGCGTTGACATCAGCTGGTCTGGTGGCCCACGCAGATAACGGTGTGCAGATCGACGGGATGTGGTTTCCGTCGGTGTATGTGTCCAGGTTAGCTGAGCTAACTGAGGCGGGCGGACACACGAAAGGTACTTCAGCGATGGCGCTGGAGCACGGCCGCTCGATCGCTTCAGACTGTGGTGCGTTCGTGCGCAATTACGTGAGAGGGGCTTCAATGCTTTTCTCGATGTCGGGGAGCGAGGCGACGGCGGAAGCTGCGCTGACAGCGTACTTTGGTGCGGCGGCCAAATCGGATGACCGCCATCTGAGGCACGCGGTGGTTTCACCCTATTATTGGGTGGAGCCGACATCGCTGCTACCGGCTGACCTGCTCGGTACGGGGGGTGAGAAGGCGGGTTGCGGCTCAGTGGTCACACCTGGGCTGACGTCGGAGCTGCCGTTGTTCACGCGGGTGAACAATTTGAGCTCAAATGGTAGTGTGCGGTCATCGTTGCAAGTTGAAATGCGGTCTGCACGCACCAGCGGGTTGGTTTGCCACCTCAATGGCAACCCGGCTAATGGGCTGGGATGTGTGAGCGTGAGACAGATGGACCCGGAGTCGATCGTGTTGCCCGGTGCTTCGGAGGGACCGGATGTCGACACGGTTCGTAAGAGGTTGTTGAAAGGGAAAGACCTGGCGGACTACCTGTGGAAGCGGGGTCAGTCACCTCTCCCGGCCCCGAGCGAGTTCCTGAATATAAAGGGGGGACTTGGTTTGGTTGTCCGCCACGAGACGTCGGAGTATGACGATTTCGTGAGCGAATTCGAGCACGTTCCCATGAACAATGAGCTCGACGGGGTGGTGACCTTTAGGGTGTCGCGGCCGCGGGGATTCTCGCTGGGGCCGGCAAACAGCGAGGATCGGGCAAGGCGTAACGCTCGGACTCGTGCTGCAGCTGCACTGAGCGCAGCAAGGTCACGTTCGAACTGGGGCGCAGCGGGGGTTTACGAGGAGATGCCGATCTCGTTCACTCCACCGGTGTTCACACGGCCGTCAACGGGCGCGGAGGTGATGCTGGACAAGGCCGCAAAACACGAGCACTTCGGAGTTTTGGCGGGTTCGACGCCTGTGTCCCGTGGCAACGAGGGCCTCGGCACACAGGGCGGAGGCGCTAGTGCCGCTGGTTCGGCGTTGCGCGCTCAAGCGGGGCACGGCGCGACGATACTGCCCCGCGTGACAACGCAGCGCGTACCGACAGGTGGAGGAGCGACGACGTCACAAGTTCCGCCGCTAGATGCAGCCATCGCGGCAAGGAGCGCTGCCCCGGGCGACGTAGGAGTCGTTCCGGAGGCGGCGCCTGAATGAGTGGATCGCGTGGAGTTGCGGCAGGGCGTGCGGCTGGGATGGGGCACTTAGGTGCATACATGCTGTCGCAACTCCCACGGGTCGGAGGAGTCTGGGGAGGCGCAGAACCAGAGAGCTTCGGGCAGCAGATTGCGGCCGTCGGGCACATGAGTAGTACTGGTGACTCATTGCACGCGGCGGCCCTCAGCATTCTGTGTTGTGAATATCCTGTACAAGTATCCTTAAAAACAAGCGAGATCATATCTCTCGCGTGGCACTCTTTTGGTGGTGACGGCAGCGCTGGCGCTTGCTTCGATTCGATCTTTAGTACCAGCAGCAAGAAAAATGGTAAACGTACAAAAAACGAGAATAACTCAAAAAACACAAAAAAGATTGAAAAAGCTGACACACAAAAAATACAGCGCTTGGTGCACATAAGCGGCGCTGGTAGGATGGTTGAAGGCAAGCGTGGCGGTAAATTGCTGAGGGCGCGGTTCTTGACGGGAGAAGCGCAGAGCAGGGGTAGGGCACGCATGGCTAGTAAGGAAGAACGCAAGCGCATCTTTCCTATTAAGCTTAGTCATCCAGGTGCAACGAATAAGGTAAACTGTTACCTTTCGGAGGTCTATCCGATGGTGGAGCAGTACTACCCACAGGTGGCTGCTGAGTTCTGCGGAGTAGTTGGTTGTGGGTTGAGCGGCGTGTACGATGATCAGGCTACAGCAATACTGATCTACGCGTGTGCGCTTAGTAGGTCTGTAGGCAACGCGGTGGAGCTCGCTTTCTGCATGGTCGCGGAGCCGGATACGGCAAAACAGCTCTCCACGACCTTAAAAGCATTAGGTTTGAATGCGACACTCGTCGGTGCGAAGCTAGTCGAAGGTAACTCTTTGTTGGGTAGAGGAGTGAATCCGGCTGACCTAAAGGCAGAAGCTGCGTACCGTTGCTCGAAGGCGGAAGCAGAGCGTGGCTGCGTGCACTTCGATGAAGATGATCTAAGAAGGGTCGTTGATCAGATCTTAGATGAGGAGATGAACGGGGAGCCGAACTTCAGGGAGCCGGAGGAACTGTGGTTGCGGCGCTGGGAGTGGTGTGTCAACGGAAGTCATGCTCAGTTACTGTCGCGCCTAAAGCCGCAGTATAAAGTGAAGGACATCCCCGGGGTCACACAGTGGTACAGGAGAATGTTTGCGGAAGCAGTGCAGGACGAGCCTATCACAGGATGGGACGGCGAAGTGGTAGTTTCAGCAAGCGAGAAGCTGGAGCATGGCAAGACGAGAGCGATCTTTGCGTGCGATAGTATCTCGTACTTCTGCTTCGAGCACTTGTTAGGACCGGTGGAAGCCGCGTGGCGTGGCAGACGGGTAGTGTTGGACCCCGGGTCGAATGGACACTCTGGTATGTGTGAGCGGATCAGAGGCCTTAGAGCCCGTGGGTCGGTTGCCCTTATGCTAGACTATGATGACTTCAATAGCCAACATACGCTGCGCAGCCAACAGGTGGTGTTGGAAGCACTGATTGAGAAGACCGGGTACGACGGCAATCTCGGTAAGAAGCTTGTCGAGTCGTTCGAACGTATGGAAATCTACGTGAATGGCTCAAGGGTTGGCAGGGCACAAGGCACTCTCATGTCCGGCCACCGTGCGACGAGTTTCCTGAATAGTGTTTTGAACGCCGCTTACATCAGGTTAGGAGTAGGAGAAGAGTTGTACGGGCGCTGCTGGAGTGTGCACGTCGGGGATGACGTGTTTATGAGTGCGGATTCATATGAGAGCGCTGCGATGGTCATGTCTCGCATGAAGCATACCGGTTGCAGGTTGAATCCGAGTAAGCAGAGCGTCGGGAGTTACTCAGCGGAGTTTTTGCGGGTAGCTCTGGCAGATGGTTTTTCGGTGGGCTACGTAGCTCGTTCCGTGGCTAGCATAGTTTCTGGCAACTGGGTATCAGAGATGAAGCTTGCCCCAATGGAAGGGCTACAGACGATGATGCAGAGCGCGAGGACACTGATCAACAGATCGCAGGATGAGGAGATCTACCGGCTGTTGTGTCCGAGCGTTAGGCGTATGACGGATATGAGTCCAGTAGTGGTAAACGGGCTTCTATCGGGGGCTGTGGCTTTAGGTGAGGGGCCGCAGTACAGGTCATCAGGGCAACGGACGGCTCTGGTAGTAAAGCCAGCTGATCTAACGAGGTTAGAGCAGGATGAGGCGGCTTGGCCTAAACACGCGACTGAAGACTTCTTGAGTCATGCGGTGTCTGACGTGGAGATCTTGGCGTTGAGCCTGACGGGCGCGTCTGTGAAGTCGGCTATGTTAACTGCTTCGTTTTCGAAAGCGGTTGCAAGTGGCAAGCAGACCCCTGAGCGGCTGACAATAAGAAGGACAGTCAATTACGCTGCGCGAGGAGTCGATACAGTGCGCGAGTTGTTAGGGCGGCGTAAGAAGCCAGGAGTGTTAGCGAACTACCCGGTTTTGCACTTAATGCGGAACCAGCTAAAACCGGAGCAGCTGCGCGCGGTAGTTCGTGAAGCTGGAGGCAACGGGAATGCGAACGACATTCTCTTGGAAGCTTGGGGAGCTGAGGCGCGCGGGGTGATAATCAGGGGTTGGTTATCATACTCAGACGCATGTCTGCTTAGCTCTCGGACGTCATCAGACGTCATAGTTGCAGACAACCCACATTATATCTAAGTAGGTAGGAACCTTCTTAGATCGATATTACACGCCCCGCAGATCGCAACAGCGTC

>Thecaphora_thlaspeos_eimeriavirus_3

TCCTACCTTATGTAGGGCAGACATAGCTTATCATACTCAAGGTAGTGGCTCCCACCACTATCGCTTGTAGTCCACTTGCGTCTCAGATAAGCCAGACCTCTGGTGTATGCTGTGTAAGCCTGGCCTCCGAAAGTCGTACCTACCGTAGCCTCAAGTGCGGGGGAAGTTCATCAGGAGACTGCCAGTTCGTTAAATCGATGGACAGTTTTGGCAGAGCATGGAGGGTGCCGGTGAAGTGCAAAACCAAATCTACAGACTACAGACAACTACTACTTGATATAAGACTACCCAACTTACAACCCAACCAACTAAGCCAACTACAATGCTGGCCTCTATGTTGAGCGCCCCTCAGGGCACATTCCTCGTGAACGTCAAAGACGCGCGCAGGTACAGGTCGGATCTCACGACCAGCACTCCTATGTTAGGAGTTAGAGACAGCTCGACCAAGAGCATCGTCTACGAGGTGGGCAGGCGCCACACTAGTGTCAAAGACGCTTTGGCCGCCGCCGACAATTCGGTTCCCGGCCCCGATATGTCCGTGAAGACCAGCTTCACCACCTCAGCTGATTTCTCTGGCCTGGCCAAGAGGTTCAGCAACTTTTCGGGCATCTGGACCCAGATGGACCTGGCTGGAATAGTCGAGCGCCTGGGCAAGGCGATCGCTACCCAGTCCCTTTTCGCCGGGGTTACCACCAAAGCTATGCGAGGTGGCAACCCCATTGACATAACCACCCTGTCGAACGTCACCCGCCCTGTGACGTCGTCACTCACTTCGGTGTTCATCCCCCGTCTCGTGGACAGCCTCAAGTCCCAGGACGTGTTCGCCGTTTTGTGCGCTGCCGCCAACGGCGAGGGAGCAACGGTCGTTACTGACTTGGTCTCCCTCGACGCGAACTCGAACTCGCCGGAGGTGCCGCAGTGTGAGGGCCAGCATTTGGCTCACGCCTGTGTCGAAGCCCTGCGCATACTAGGCGCAAACTACCACGTGAACAACGCGAGCGACGTTTTCGCCTACGCGCTCACCCGTGGTATCCATTCGGTGGCCAGTGTCGTCGGCCACACTGACGAAGGAGCCTACCTTAGGAAGGTGCTCCGCACTACTGGGTTCGCGACGCCCTACGGTGGAATCCACTACTCCACCAACAACTTCCCCGGTCTGCCGGCACTTGCGGCAAAGACCGACAAGGCCATAGCCGGGTGGGTCGACACGCTCGCGCTGGCTACCGCGGCCATCGCTGCACACTGCGACCCCTTGGTCGAGAGCAACGGCTCGCTGTTCCCGTCGGTCTTCGTCTCTCCTATCGGTGAGTACGAAGAGGCCGGGGGAGCGGGCGAGGGTACAACCGACGATGCCCTCGCCCTTGCCCGCGAGATTGCAGCTGATGCACCTCGCTTCGTCGGAAACTATGTGGCAGGCCTCACGAAGATTTTCGAGGCCATCCCCAGCGGTGACTCTCTCCCGGCCCGCCACCTCGAGAACTGCTTCATCGTTGCCTCGAAGACGGCTGACCGTCACTTGCAGCGCAAGACCGTTGCCCCCTACTACTGGATCGAGCCCACTTCGATCGTGAGTGCCAACTACCTCGGCACGAAAGCCGAAGAGCACGGCTACGCTTCTCTCGCGTCTCCAGGGTGCGACTCGGTGATGAAGGGTTTCTCGTACATCGAGGAGAAGACGCACGGTTCGAGTGATGCTGGGAGTTACGTCGTGGAGATGAGGAGTGCCCGTTCCTCGGGGCTGGTCGTACACCTCAACGACCACCCCCTCAACGGACTAGGAGCTCTTAAAGTCCTTCAGATGGACCCTACGGGCGTAGTAATGCCAGGCTTCCGGTCCGGGGAAACCGCCGACGTTGCCACCCGGCTCCGTGACAACGAGCCGATCTCCTCGTATTTGTGGAAACGCGGCCTTAGCTGTATACCGGCACCGGCCGAATTCCTTAATACAGGGATAACTCTCGGTGTGTCGGCCGTCCACATGAAGTGGGATAGCCTGTACAGACATGCCGAACCAGAACACCTCCCCGACCCCCGCGAGATGTTGGACTGCGATGTGACTATACGGGTGTCACGCCCCCGAGGGATCGGTACTGCCCCTAGCAACTTCGAAGGAAGCGATAGGAGCAGTGCGCGGTCGCGGGCGTCCATGGCCCTCAGTATGGCCAGGGCGAGGTCGCGGTACTACCGCGACACGGGGGCGGAGGCGATGATGACCAGCGCGACCGCCCCCCGGGTTCGTAAGAAAGGGGAGGTCGCGACGGTCGTGCTACCTGGAGCGTCAGTAGATGACGTGAAAGCGATGCTCGACCACACGACCTACCGGGGGGCGGCGGCTGACAAGGGCCCTGAGGTCACGGAGTATCGCACAGATAACAACGTGACGCATCGGGGCCCTGTCGCCGGCGTGAAGCTGCACCTTTCCGATGACCGCGGACCCCGGGCCAAGAGTTCTATACCTGGCTCGGGGTCCGGTGGGTCCGGATCGGCGGTGGAGCAGTTGGCACGGAGCGAGGCGGCTCCCCGGGAGGACGGGAGGACACTTAGTACGCCGGAGGGTAGCGGGCTCGCGCCCCCTAACCCCGAGGCGGCCGCTCAATGAGCGCTGAGCGTCTCGCCGAGAGGGTCGCTCAGCTGGGCCTCCTCGGCGTAGCTCTCTTAAGAGAGCTCGACAATCAGCTGTTCCTCCAGTTTGACTCTCTCACTACCGGTGAGCAGATCAACTGGATCGGCCAGGTCACGTTATCCAGAGGGGAGCGTGCAGCCTGTGCGTGTAGTGTGTTGCTGTGTGACTTCCCGGTACAGGTCGGGATCTCGGATAAGGCCGTGATGTGGCTTGTGCACTCGGCAATTGCGGGCCTGGACCCGGGGAAGAAGGGTGCGGGGCTACCTCCGGAGCCCGGTATGCAGGGGTACCTCAGAGAAGGGTTCAAGCTCAAGAGCCACCCTGGGGCTACTAACAAGGTTAACCTGTACGCGTCGGAGGTGCTCGACGGGATATTCGAGGCCGACGCACTATCGTACAGATTTTGCCTGAGGCCCCTCCCGTACTTATACGGGCGTGTGTTCAACGATCAGTTAACTGCTTACCTGATCCACGGCTACGCACTTGAACGAGCGCGAGTGGAGGACGCCTACTTCTTGGCAGCCCGTATGATCCTGGACCCGGCCGGATGTAAGTCCCTCAGCGGTACCCTCAAAGCTCTGGGGGCCAATGCAGGGCGTTTGGGCTCCCTGTTCGTGGAGTGTGATACACTTCAGGGACGTGGGGTAAGACCCGTCAACCTCGCAGCAGAATGCGAGCGCAGGTGCGACCCAGTGGCGTGTAGCCGAGACAACGCTAAGTTCGATCGTGACGCGCTCACTGCTATGGTGCGTACTATCCTTGACGAGGAGCTGGGGGGCAGGACCATCGAGCATGAGAAACTCGACAGTTTCTGGGACGGGCGCTGGGCCTGGTGTGTGAACGGCAGCCACTCTAAGCTGCTACAGCGCAACAAGCCACAGTATAAAGTCTCGGAGATCCCCGGCGTCTCACAATGGTACAGGAGGATGTTTGCGGAAAGTACCGAGAACGAACCAATCAGTGGATGGGATGGTTCGGTCTACGTCACAGCCTCAGAGAAGCTCGAGCACGGCAAAACCAGGGCTATCCTGGCTTGTGACACGGTAAGTTACTTTGCGTTCGAGCACCTCCTCAGGCCGGTGGAGAAGGCTTGGCACGGGCGGCGGGTGCTGCTGGACCCGGGGTCCTCCGGCCACGCAGGTATCGTTGGCAAGGTCAAGGAACTGAGGGCGAGGGGGGCCGTCTCGGTGATGCTGGATTTTGACGACTTCAACTCCCAGCATACCCTGGAGGCCCAGAAAATCGTGATCCGAGAGGTGTGTGAGAGGGCCCAGTACCCCGCAGAACTAGCGGACAAGCTTGTGGGGTCCTTCGACAGAATGAACATCTACCACTCGGATCGCCTAGTAGGCACAGCAGCTGGCACGCTTATGTCAGGACACCGCGGGACCACATTCCTTAATTCTGTCCTTAACGCTGCGTACATAAGGCTAAGCCTGCCGGCGGGTGTCTACGAAAGGTGCTGGTCGCTGCACGTGGGAGACGACGTCTTTATGAGCGTCCCTACTTTCCACGACGCTGAAGCGACGATCCGTGGTATGAGTTTATCTTCGTGTCGCTTGAACCCCATCAAGCAGAGCGTGGGTACGGTAACTGCCGAGTTCCTAAGAGTAGCGGTAACGGAATGGTCGGCACAGGGTTATCTCGCAAGGAGCATCAGTTCGGCGGTGAGCGGAAACTGGGTGAACGAGACAAGTCTCTCCCCGCGAGAGGGATTGTCCACGATGATCCAGAGCAGCCGGACCATGTTTAACCGATCAAGAGGTTGCGATTTTTCCGTCCTCTTGGCTAGGTCGTGCGCCAGGATGGCCGGTAACTCTCAGAAGAATGTTGTGCGCCTGCTGCGAGGCGAAGTATCGCTCGGTGCGGGCCCCGTGTTCGGGAATAAAGGGCTGTACCGAGAAGCCCGGCTATCTGTACGACAGCCGGACGACGACAGACTGCCGCCAGGACTACCCCACCTGGCGACGAGTGCCTACCTGAGTCGGAAGCTGCAACCGCCTGAATTGTACGCGTTAGAGAGACTGGGACACAGTGTTACGTCGGCGATGATAAGGTCTTCTTTTCAGAAGGCTCTATTACGTGGCACAGACTGTGAACCCCAGCTCGACATCCGTACCCTTAAGGCGGAGGTGCCTATCGGCTCGACGAGCTTGGAGGTCGCGCTGAAGTTCCGCCCACCCCCCGGAGTACTGTCTGGATATCCAGTCTTGCAACTCATGAGGGAGGGGCTTAGCAGGAGCGAAGTGCGGGAGCTCGTGAAGGTGGCTGGCGGGGACGCCAACGCCCCCAACATACTGTTGGAGGCTTGGGGCCCCGAGAGCGTCGGGTGTGTCATAGAGGGCTTCGTGTCCTACTCTGACGCACGCGCGGCGTGTAAGAGGGTGACCATAGGGGTCATCTACACCATTTACGCTTATTACATCTAAGTGTAAATGAACACCCACTCAGTGGGAC

Color code: satar codon; sotp codon
